# Energy efficient network activity from disparate circuit parameters

**DOI:** 10.1101/2021.07.30.454484

**Authors:** Michael Deistler, Jakob H. Macke, Pedro J. Gonçalves

## Abstract

Neural circuits can produce similar activity patterns from vastly different combinations of channel and synaptic conductances. These conductances are tuned for specific activity patterns but might also reflect additional constraints, such as metabolic cost or robustness to perturbations. How do such constraints influence the range of permissible conductances? Here, we investigate how metabolic cost affects the parameters of neural circuits with similar activity in a model of the pyloric network of the crab *Cancer borealis*. We use a novel machine learning method to identify a range of network models that can generate activity patterns matching experimental data, and find that neural circuits can consume largely different amounts of energy despite similar circuit activity. Furthermore, a reduced but still significant range of circuit parameters gives rise to energy-efficient circuits. We then examine the space of parameters of energy-efficient circuits and identify potential tuning strategies for low metabolic cost. Finally, we investigate the interaction between metabolic cost and temperature robustness. We show that metabolic cost can vary across temperatures, but that robustness to temperature changes does not necessarily incur an increased metabolic cost. Our analyses show that, despite metabolic efficiency and temperature robustness constraining circuit parameters, neural systems can generate functional, efficient, and robust network activity with widely disparate sets of conductances.

## Introduction

Neural activity arises from the interplay of mechanisms at multiple levels, including single-neuron and network mechanisms. Several experimental and theoretical studies have found that neural systems can produce similar activity from vastly different membrane and synaptic conductances [1–6], a property sometimes referred to as parameter degeneracy [7, 8]. Such parameter degeneracy has been argued to be a prerequisite for natural selection [7] and translates into potential mechanisms of compensation for perturbations of the systems’ parameters [3, 5, 9–14]. However, in addition to a specific target activity, neural systems are likely subject to additional constraints such as the requirement to be energy efficient [15–17]. In order to understand experimentally observed variability and probe potential compensation mechanisms in functioning neural systems, it is thus crucial to characterise the extent of the systems’ parameter degeneracy under such additional constraints.

Neuronal activity accounts for the majority of the energy consumed by the brain [18–20]. Energy is stored in the ionic gradients across the cell membrane, and consumed mostly by action potentials and synaptic mechanisms. Maintaining the ionic gradients requires the action of ion pumps, which consume ATP [15, 21]. Previous work has investigated the metabolic efficiency in small neural systems, often at the single neuron level and with few ion channels (often sodium, potassium, and leak) [15, 22, 23]. In these studies, it has been demonstrated that energy consumption of single neurons can be reduced by tuning maximal conductances or time constants of gating variables, while maintaining electrophysiological characteristics, e.g. spike width. However, questions regarding energy efficiency of neural systems remain: First, it is unclear whether previous findings in single neurons [24–26] extrapolate to neural circuits with a large diversity of membrane and synaptic currents [12, 21, 27]. Second, the question of how strongly metabolic constraints impact parameter degeneracy remains unaddressed: Are energy efficient solutions confined in parameter space or can disparate network parameters generate energy efficient activity? Lastly, metabolic cost is only one of many constraints under which neural circuits operate, and it is often unknown whether energy efficiency trades-off with other constraints (for a study of how energy efficiency trades off with temperature robustness in a single neuron model of the grasshopper, see Roemschied et al. [28]).

Here, we investigate how energy efficiency constrains the parameter degeneracy in the pyloric network in the stomatogastric ganglion (STG) of the crab *Cancer borealis* [29, 30], a canonical example of a neural system with parameterdegeneracy [5]. The pyloric network produces a triphasic motor pattern, and consists of a pacemaker kernel (anterior burster neuron, AB, and two pyloric dilator neurons, PD), as well as two types of follower neurons (a single lateral pyloric, LP, and several pyloric, PY, neurons), interconnected by inhibitory synapses. A model of this circuit with three model neurons (AB/PD, LP, PY), each with eight membrane currents, and seven inhibitory synapses (Fig. 1 a, details in Methods) has been shown to be capable of producing similar network activity with widely different parameters [5].

**Figure 1.**
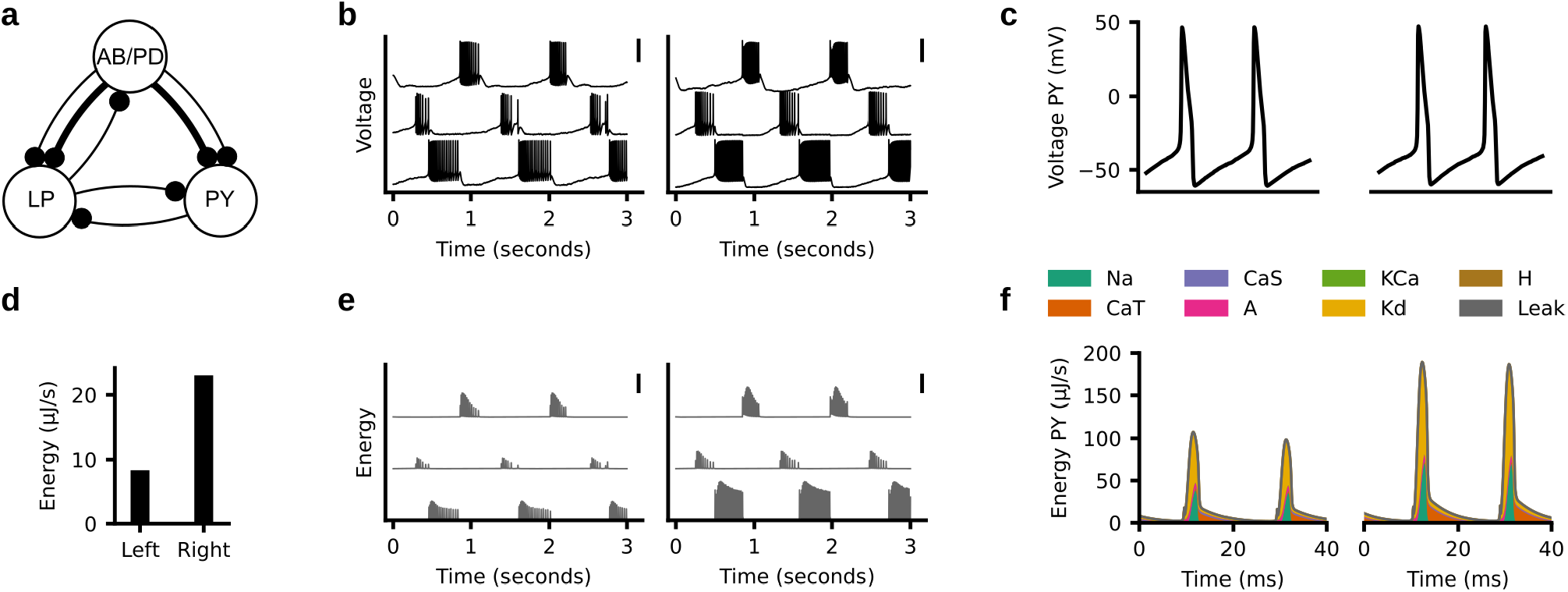
Similar activity with different energy consumption. (a) Computational model of the pyloric network consisting of three model neurons (AB/PD, LP, PY) and seven synapses. (b) Two model configurations with similar circuit activity (traces from top to bottom: AB/PD, LP, PY) despite different circuit parameters (parameter values not plotted). Scale bars indicate 50 mV. (c) Close-up of two spikes in the PY neural activity shown in (b). (d) Total energy consumption divided by the duration of the simulation (10 seconds) for the traces shown in (b). The left circuit has 3-fold lower metabolic cost than the right circuit. (e) Consumed energy at each time point. Scale bar indicates 100 *μ*J/s. (f) Energy consumed by each of the ion currents during the two spikes shown in (c).

We start by characterising the parameter degeneracy of this model: We apply a recently introduced machine learning tool for simulation-based inference, Sequential Neural Posterior Estimation (SNPE) [14] to estimate the full set of membrane and synaptic conductances for which the model reproduces experimentally measured electrophysiological activity. We reduce the number of model simulations required to run SNPE by introducing an additional classifier which detects and rejects parameter-combinations that produce non-bursting model outputs [31]. After characterising the parameter degeneracy in the model, we show that disparate circuit configurations can have different energy consumption despite similar activity. However, a significant parameter degeneracy is present in the model even when enforcing circuits to have *both* similar activity and low energy consumption. Furthermore, energy consumption is linearly predictable from circuit parameters, allowing us to identify tuning mechanisms for low metabolic cost. We then show that individual neurons in the pyloric network can be tuned separately to minimize their energy consumption, and thereby achieve low energy consumption at the circuit level. Finally, since the crab *Cancer borealis* is subject to daily and seasonal fluctuations in temperature, we study the trade-off between metabolic cost and robustness to changes in temperature [32–35]. We find that metabolic cost can vary across temperatures, but that the pyloric network can produce functional, energy efficient, and temperature robust activity with disparate parameters.

## Results

### Disparate energy consumption despite similar network activity

We studied the metabolic cost in a model of the pyloric network (Fig. 1a). In this model, disparate sets of maximal membrane and synaptic conductances can give rise to similar network activity [5]. As an example, we simulated two such circuit configurations (Fig. 1 b) and computed their metabolic cost using a previously described measure of energy consumption [36]. In this measure, the energy for each ion channel is the time integral of the product of the membrane current and the respective difference between the membrane voltage and the reversal potential. The energy consumed by the entire neural circuit is the sum of the energies across channels of all neurons (details in Methods).

Although the two simulated circuit configurations produce similar network activity, even at the single-spike level (Fig. 1 c), the total energy consumption (Fig. 1 d) as well as the moment by moment energy consumption differ substantially (Fig. 1 e). A closer inspection of the energy consumed by each current in the PY neuron during the action potentials [37] shows that the difference in energy between these two network configurations is also evident in the energy consumed by the sodium current Na, the delayed-rectifier potassium current Kd, and the transient calcium current CaT (Fig. 1f).

### Disparity in energy consumption in models matching experimental data

The example above illustrates that the model of the pyloric network can, in principle, produce the same activity with different metabolic costs. However, it is unclear how broad the range of metabolic costs associated with the same network output is. In order to address this, we need to identify the full space of maximal membrane and synaptic conductances (31 parameters in total) that match experimental measurements of network activity and to characterise the energy consumption of each of these configurations.

We used a recently introduced machine learning tool for simulation-based inference, Sequential Neural Posterior Estimation (SNPE) [14], to estimate the set of circuit parameters (the posterior distribution) consistent with data and prior assumptions about the parameters. In SNPE, parameters which specify network configurations are initially sampled from the prior distribution (in our case a uniform distribution within plausible parameter ranges) and used to simulate network activity. Subsequently, a neural-network based density estimator is trained on these simulated network activities to learn which parameter sets produce network activity that is compatible with empirical observations. In order to generate the training data for the neural network, SNPE requires millions of model simulations to accurately infer the set of data-compatible parameters. To improve the simulation efficiency and make the neural network predict parameter sets that more closely match experimental data, we introduced a modification of the algorithm (Fig. 2a). Specifically, a technical challenge for SNPE is that parameter sets sampled from the prior distribution might produce simulation results that are not ‘valid’, i.e. produce clearly non-sensible data: E.g., if there are no bursts, phase gaps between bursts are not defined (Fig. 2a, forth panel, red). For SNPE, these ‘invalid’ simulations are discarded immediately. In order to reduce the fraction of simulations that are discarded, we introduce a classifier to predict whether a parameter set will lead to a ‘valid’ or an ‘invalid’ simulation output [31] (Fig. 2a, second panel). Once the classifier is trained on an initial set of simulations, parameters are immediately discarded without running the simulation, if the classifier confidently predicts that the simulation will be invalid (details in Methods). We name the distribution of parameters that are accepted by the classifier the ‘restricted prior’ (Fig. 2a, third panel). Once sufficiently many valid simulations are performed, SNPE proceeds by training a deep neural density estimator to estimate the posterior distribution over parameters of the model [14] (Fig. 2a, last two panels, proof of convergence to the correct posterior distribution in Methods).

**Figure 2.**
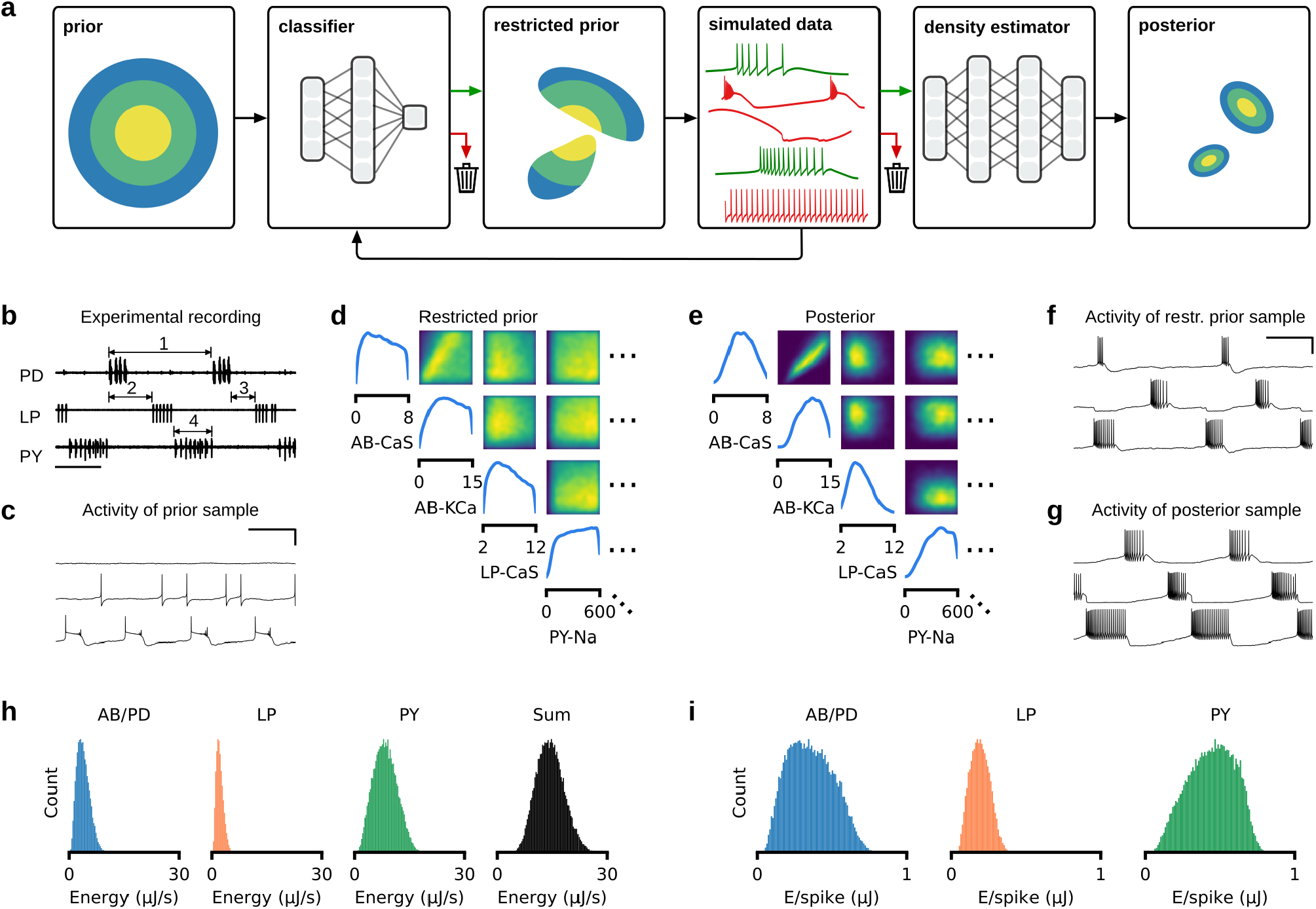
Bayesian inference reveals wide range of energy consumption. (a) Inferring the posterior distribution by combining a rejection classifier and a deep neural density estimator. First, a classifier (trained on an initial set of simulations) predicts which circuit parameters sampled from the prior produce ‘valid’ simulation outputs. We then proceed by sampling from the part of the parameter space that is accepted by the classifier, i.e. the ‘restricted prior. All ‘valid’ data (green) are used to train a deep neural density estimator, all ‘invalid’ data are discarded (red) [14]. Once this estimator is trained, it can be evaluated on experimental data to return the posterior distribution over model parameters. (b) Experimental data recorded from the pyloric network [38]. Arrows indicate a subset of the physiologically relevant features, namely the cycle period (1), phase delays (2), phase gaps (3), and burst durations (4) (see Methods for details). (c) Simulation output from a parameter set sampled from the prior distribution. The traces are: AB/PD (top), LP (middle), PY (bottom). Scale bars correspond to 500 ms and 50 mV. (d) Subset of the marginals and pairwise marginals of the 31-dimensional restricted prior, i.e. the subspace of parameters for which the model produces bursting activity. All maximal conductances are given in mS/cm^2^. (e) Subset of the marginals and pairwise marginals of the posterior distribution, i.e. the subspace of parameters for which the model matches experimental data shown in panel (b) (full posterior distribution in Appendix 1 Fig. 1). (f) Sample from the restricted prior producing bursting activity but not matching experimental data. (g) Sample from the posterior distribution closely matching features of the experimental data. (h) Histograms over energy consumed by each neuron (blue, orange, green) as well as by entire circuit (black). Trace with lowest energy consumes 9 times less energy than trace with highest energy. (i) Same as in (h), but for energy per spike.

We used this procedure to infer the posterior distribution over maximal membrane and synaptic conductances of the model of the pyloric network given salient and physiologically relevant features of experimentally measured data. These features are the cycle period, burst durations, duty cycles, phase gaps, and phase delays of the triphasic rhythm (Fig. 2b, details in Methods) [38]. As in previous studies [4, 5], we did not constrain the model of the pyloric network by the number of spikes or the spike shapes. Below, we describe the results obtained for a specific experimental preparation. We qualitatively reproduced all results with two additional experimental preparations (Appendix 1 Fig. 11, Appendix 1 Fig. 12, Appendix 1 Fig. 13, Appendix 1 Fig. 14, Appendix 1 Fig. 15, Appendix 1 Fig. 16) [38].

When simulating the pyloric network model with parameter sets sampled from the prior distribution, 99% of simulations do not produce spikes or bursts and hence characteristic summary features of the circuits are not defined (Fig. 2c). The restricted prior (Fig. 2d) is narrower than the prior distribution, but considerably broader than the posterior (Fig. 2e, full posterior distribution in Appendix 1 Fig. 1; comparison between prior, restricted prior, and posterior in Appendix 1 Fig. 2). Parameters sampled from the restricted prior often produce activity with well-defined summary features (Fig. 2f), but do not generally match experimental data, whereas samples from the posterior closely match experimental data (Fig. 2g). By using the classifier to reject ‘invalid’ simulations, we required half as many simulations compared to ‘classical’ SNPE [14] and achieved a higher accuracy (Appendix 1 Fig. 3). For the subsequent analyses, we only considered posterior samples whose activity was within a prescribed distance to the experimental data, and discarded all other samples(details in Methods). We simulated 1 million parameter configurations sampled from the posterior, out of which approximately 3.5% fulfilled the distance criterion, leading to a database of 35,939 parameter sets whose activity closely matched experimental data. Sampling from the prior distribution rather than the posterior would have required approximately 600 billion simulations to obtain 35,939 parameter sets that fulfill our criterion (60,000 times more than with our method).

We computed the energy consumption of each of the 35, 939 circuit activities (Fig. 2h). The circuit configuration with lowest total energy consumes nine times less energy than the circuit configuration with highest total energy. To ensure that the difference in energy does not only stem from different numbers of spikes within a burst, we also computed the average energy consumed during a spike (energy per spike) in each of the neurons (Fig. 2i). As with total energy, energy per spike strongly varies across parameter configurations. These results show that, despite similar circuit function, different parameter sets can have vastly different energy consumption. Below, we investigate the mechanisms giving rise to this phenomenon.

### Metabolic constraints on individual circuit parameter ranges

How strongly does enforcing low energy consumption constrain the permissible ranges of circuit parameters? We inspected the circuit parameters of the 2% most and least efficient configurations within our database of 35,939 model configurations (Fig. 3a, left). For some circuit parameters, the range of values producing efficient activity is clearly different from the range of values producing energetically costly activity (e.g. the maximal conductance of the transient calcium current in the PY neuron, Fig. 3a, middle). For other parameters, the range does not change (e.g. the maximal conductance of the delayed-rectifier potassium current in the AB/PD neuron, Fig. 3a, right). To quantify how strongly low energy consumption constrains parameters, we compared the parameter standard deviation across all 35,939 model configurations to that of the most efficient 2% (Fig. 3b,c). Most parameters in the circuit barely get constrained by energy consumption (values close to one in Fig. 3b,c). The parameters that get constrained the most by enforcing low energy consumption are the Na and CaT conductances of the AB/PD neuron, the CaS conductance of the LP neuron, and the Na, CaT, CaS, and leak conductances of the PY neuron. However, for all of these parameters, a large fraction of variability remains.

**Figure 3.**
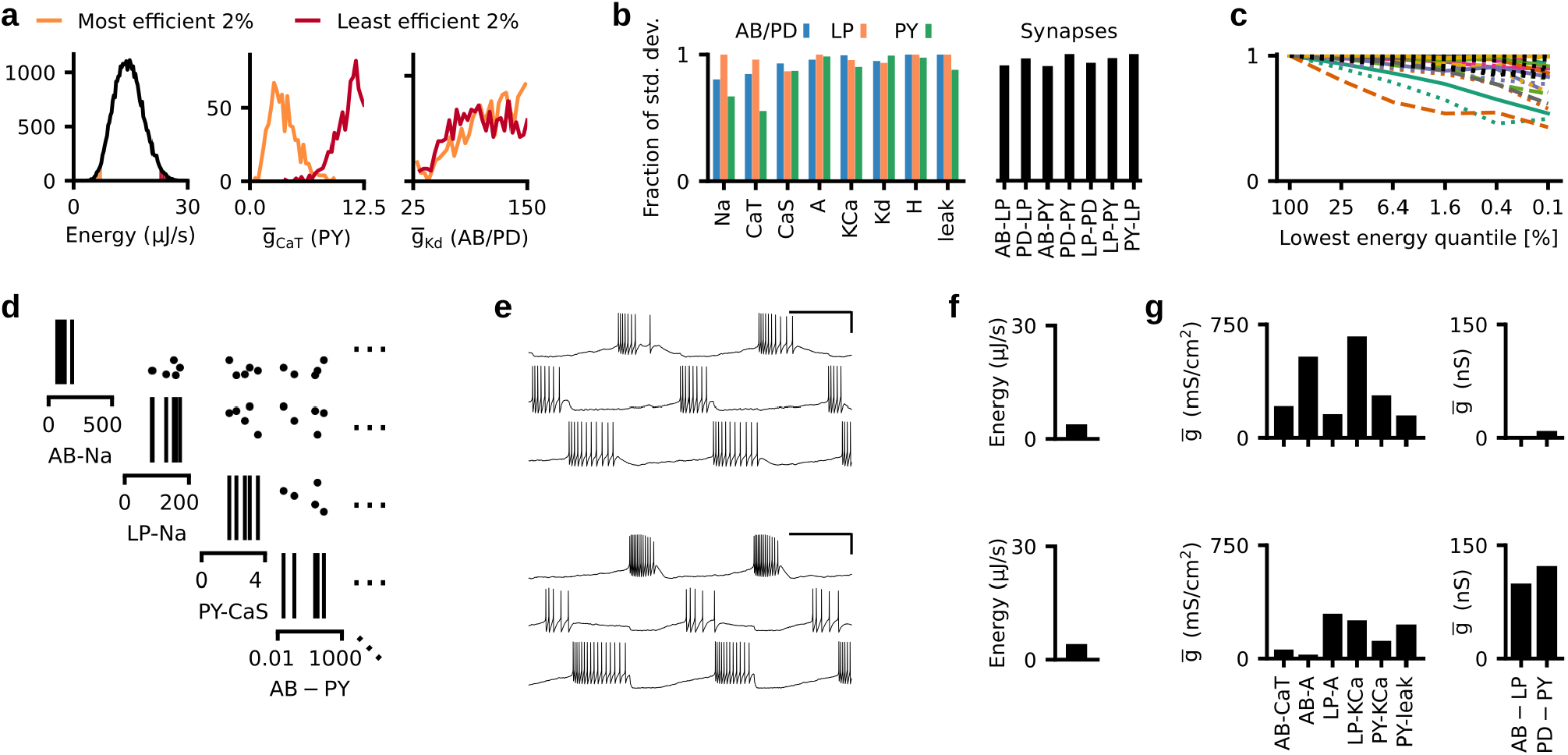
Metabolic constraints on individual circuit parameters. (a) Left: Energy consumption of 35,939 models that match experimental data. The orange area corresponds to the energy consumption in the lowest 2% quantile-red area to the top 98% quantile. Middle: Distribution of the maximal conductance of the transient calcium channel (CaT) in the PY neuron in the 2% (orange) and 98% quantile (red). Right: Distribution of the maximal conductance of the delayed-rectifier potassium channel (Kd) in the AB/PD neuron in the 2% (orange) and 98% quantile (red). (b) Standard deviation of parameters for models with energy consumption in the lowest 2% quantile. Standard deviation is normalized to the standard deviation of the parameters across all 35, 939 models in our database. (c) Same as panel (b), but for a range of quantiles. Solid: AB/PD, dotted: LP, dashed: PY. Colors are the same as in Fig. 1f. Synapses in black. (d) Subset of the parameter values of the five most efficient circuit configurations in our database. (e) The network activity produced by two of these five configurations. Scale bar indicates 500 ms and 50 mV. (f) The energy consumption of the two configurations shown in panel (d). (g) Subset of circuit parameters of the two solutions shown in panel (b). Despite similar network activity and low energy consumption, several parameters differ by more than 2-fold. The membrane conductances are scaled by the following factors (left to right): 100,10,10,100,100,10000.

In order to ensure that the remaining variability of circuit parameters does not stem from the remaining variability of energy consumptions within the lowest 2% quantile, we inspected the five most efficient configurations in our database of 35,939 model configurations. Even these five circuit configurations have strongly disparate circuit parameters (Fig. 3d). Despite having similar activity (Fig. 3e) and very low (and similar) metabolic cost (Fig. 3f), their circuit parameters are disparate (Fig. 3g). These results demonstrate that metabolic efficiency constrains the range of some circuit parameters, but it is possible to achieve low metabolic cost and similar network activity with widely disparate circuit parameters.

### Energy consumption can be linearly predicted from circuit parameters

We wanted to understand how each circuit parameter affects energy consumption. We performed a linear regression from circuit parameters (taken from our database of 35,939 model configurations) onto the energy consumption of these circuits (Fig. 4a). This linear regression achieved a high accuracy, demonstrating that energy consumption can be linearly predicted from circuit parameters (Fig. 4b; a non-linear regression with a neural network leads to similar results and is shown in Appendix 1 Fig. 5; details in Methods). The regression weights *w* indicate how strongly energy consumption is correlated with each parameter value (Fig. 4c). The maximal sodium conductance 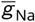 and the transient calcium conductance 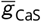 of the AB/PD and PY neuron as well as the slow calcium conductance 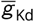 of the AB/PD, LP, and PY neuron are most strongly correlated with energy consumption: Increases of these conductances are associated with an increase in energy consumption, and thus, small conductance values correspond to metabolically more efficient solutions. The synaptic conductances are weakly correlated with energy consumption, which can be explained by the low values of the maximal synaptic conductances: The synaptic strengths range up to 1000 nS, whereas the membrane conductances can range up to 0.4 mS (i.e. 4 · 10^5^ nS), such that synapses consume only 0.08% of the total energy in the circuit. These results demonstrate that energy consumption can be linearly predicted from circuit parameters, and that energy consumption is most strongly correlated with the maximal conductances of sodium as well as slow and transient calcium.

**Figure 4.**
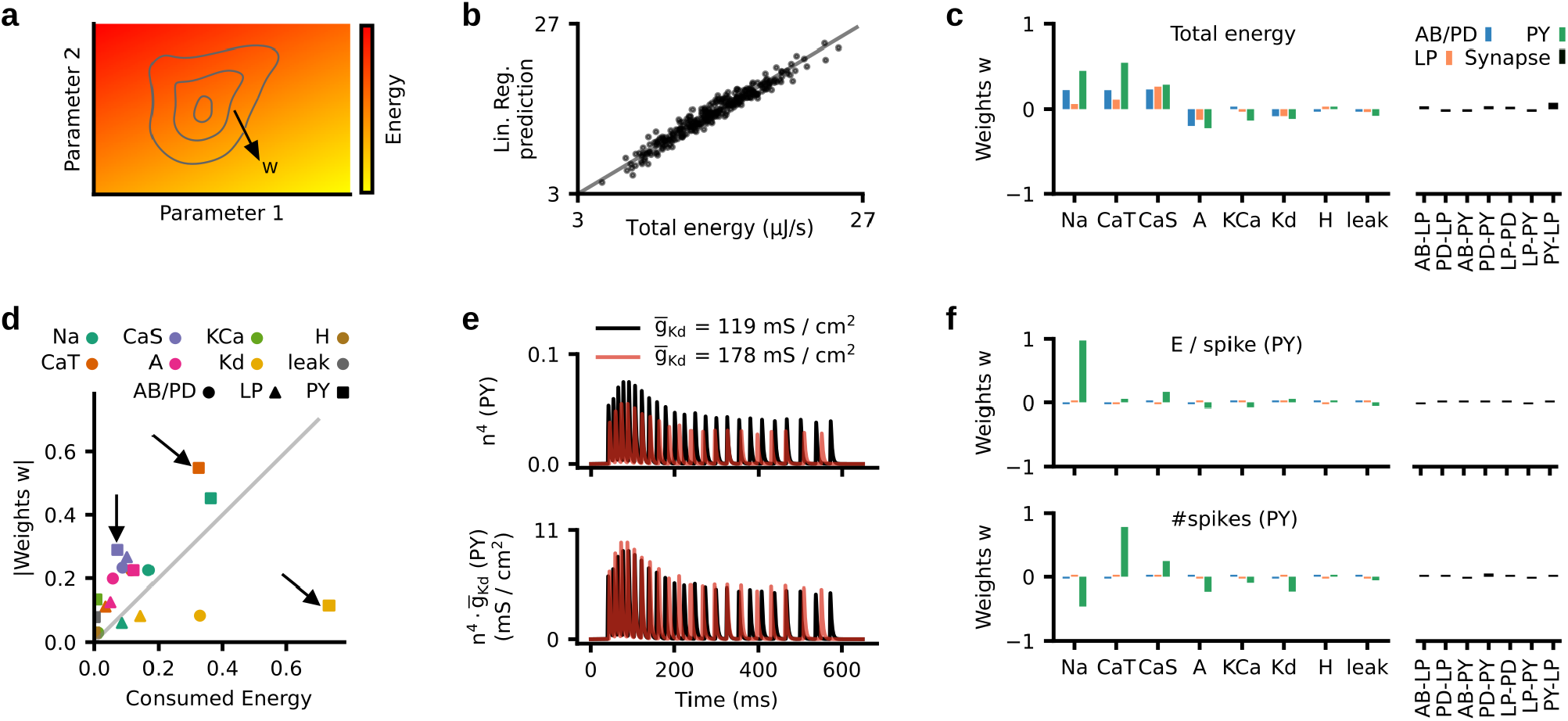
Influence of circuit parameters on energy consumption. (a) Illustration of the energy landscape under functional constraints (matching experimental activity). The linear regression weights correspond to the direction along which energy varies. (b) The linear regression accurately predicts the energy consumption on a test set of 300 circuit configurations (black dots). Grey line is the identity function. (c) Weights *w* of the linear regression. Left: Weights of the maximal membrane conductances. Right: Weights of the maximal synaptic conductances. (d) Weights *w* as a function of energy consumption (both normalized), for all membrane currents (arrows highlight three illustrative examples). Membrane conductances on the top left consume little energy, but their maximal conductances correlate strongly with energy consumption. Conductances on the bottom right consume a lot of energy, but their maximal conductances correlate weakly with energy consumption. (e) Top: The gating variable *n*^4^ of the Kd current in the PY neuron during activity produced by two circuit configurations (black and red) which are identical apart from the magnitude of 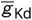. Bottom: The product of gating variable and maximal conductance 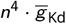 for the same configurations. (f) Top: Weights of a linear regression onto the energy per spike in the PY neuron. Bottom: Weights of a linear regression onto the number of spikes in the PY neuron.

How do different currents affect total energy consumption? Do they directly consume energy, or do they trigger processes that then require energy? We addressed these questions by comparing the fraction of energy consumed by each current (as defined by our measure of energy [36], Fig. 1 f) to the linear regression weight *w* associated with its maximal conductance (Fig. 4d). We found that some currents consume a lot of energy, although their maximal conductances barely correlate with energy consumption, e.g. the Kd current in the PY neuron (Fig. 4d, bottom right arrow), while other currents consume little energy, but nonetheless their maximal conductances are correlated with energy consumption, e.g. the CaS and CaT currents of the PY neuron (Fig. 4d, top left arrows).

We investigated the neuronal mechanisms that give rise to these behaviors. First, to understand how currents can consume large amounts of energy despite their maximal conductance only weakly correlating with energy, we investigated the effects of the delayed-rectifier potassium conductance 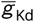 on circuit activity. We simulated two circuit configurations, identical apart from the magnitude of 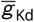 in the PY model neuron. In the configuration with higher 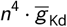, the gating variable *n* did not reach as high values as for the other configuration, thus leading to a similar effective conductance 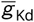 (Fig. 4e). This demonstrates that changes in the maximal conductance 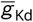 only weakly influence the current and thereby the energy consumption. Thus, despite the potassium current consuming a lot of energy due to a large flow of ions (compared to other channels), its maximal conductance 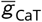 only weakly correlates with energy consumption. Second, to understand how maximal conductances can correlate with energy consumption despite their channels consuming little energy, we disentangled the correlation of circuit parameters with energy consumption into two parts: The energy *per spike* and the number of spikes. We fitted two additional linear regression models: One regression from circuit parameters onto number of spikes in the PY neuron and one regression from circuit parameters onto energy per spike in the PY neuron. We again found good predictive performance of these models, showing that the energy per spike and the number of spikes can also be linearly predicted from circuit parameters (regression performance in Appendix 1 Fig. 6). The energy per spike is strongly correlated with the sodium conductance (Fig. 4f, top), whereas the number of spikes is most strongly correlated with the maximal conductance of transient calcium (also with sodium, slow calcium, and transient potassium conductances, Fig. 4f, bottom). This demonstrates that increases in the maximal conductance of transient calcium lead to a higher number of spikes, which involve increased energy consumption through other currents. We verified this hypothesis by simulating two configurations that were identical apart from the magnitude of 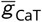 in the PY model neuron and found that the configuration with higher 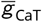 indeed produced more spikes per burst (Appendix 1 Fig. 7). This shows that, despite the calcium channel consuming little energy itself, increasing 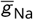 can lead to higher energy consumption by increasing the number spikes, which involve energy consumption through other currents (mostly sodium and potassium). Overall, our analyses demonstrate that currents which consume a lot of energy are not necessarily the ones that influence energy the most.

### Minimal tuning mechanisms for low energy consumption

We identified circuit parameters that correlate with energy consumption, but this does not yet address the question of which changes of these parameters will lead to the reduction of energy consumption: First, a correlation between parameter values and energy consumption does not imply a causal connection between these. Second, parameters that correlate strongly with energy consumption might have to be finely tuned to match the pyloric rhythm, thus not constituting a feasible substrate for reducing energy consumption. Therefore, we went beyond the previous analysis to investigate potential tuning mechanisms involving single and pairs of parameters that would reduce energy consumption while maintaining the pyloric rhythm.

We investigated how strongly energy consumption could be reduced by mechanisms that involve a single parameter. For instance, we kept all parameters but the maximal sodium conductance of the AB/PD neuron 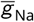 constant and varied 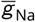 on a grid. We then estimated the energy consumption of each configuration with the previously identified linear model (Fig. 4). The energy consumption of the circuit increases with 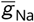. However, for too low (or too high) 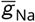, the network activity does not match experimental data (we rejected parameters for which the posterior density is too low, see Methods). Thus, despite 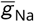 strongly correlating with energy consumption (Fig. 4c), energy consumption can be reduced only modestly when tuning 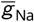 and keeping all other parameters constant.

We then investigated whether pairwise mechanisms could lead to larger savings in energy consumption. For instance, we kept all parameters but 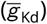 and the delayed-rectifier potassium conductance of the AB/PD neuron 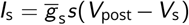 constant and varied the remaining two parameters on a grid. We estimated the energy consumption of any configuration on this grid and found that the most efficient parameter configuration is 23% more efficient than the most wasteful configuration (Fig. 5b). This reduction in energy consumption could be achieved through a simple pairwise mechanism: A reduction of sodium combined with an increase of potassium allows the network to maintain its activity (Fig. 5c,d), while reducing the metabolic cost (Fig. 5b).

**Figure 5.**
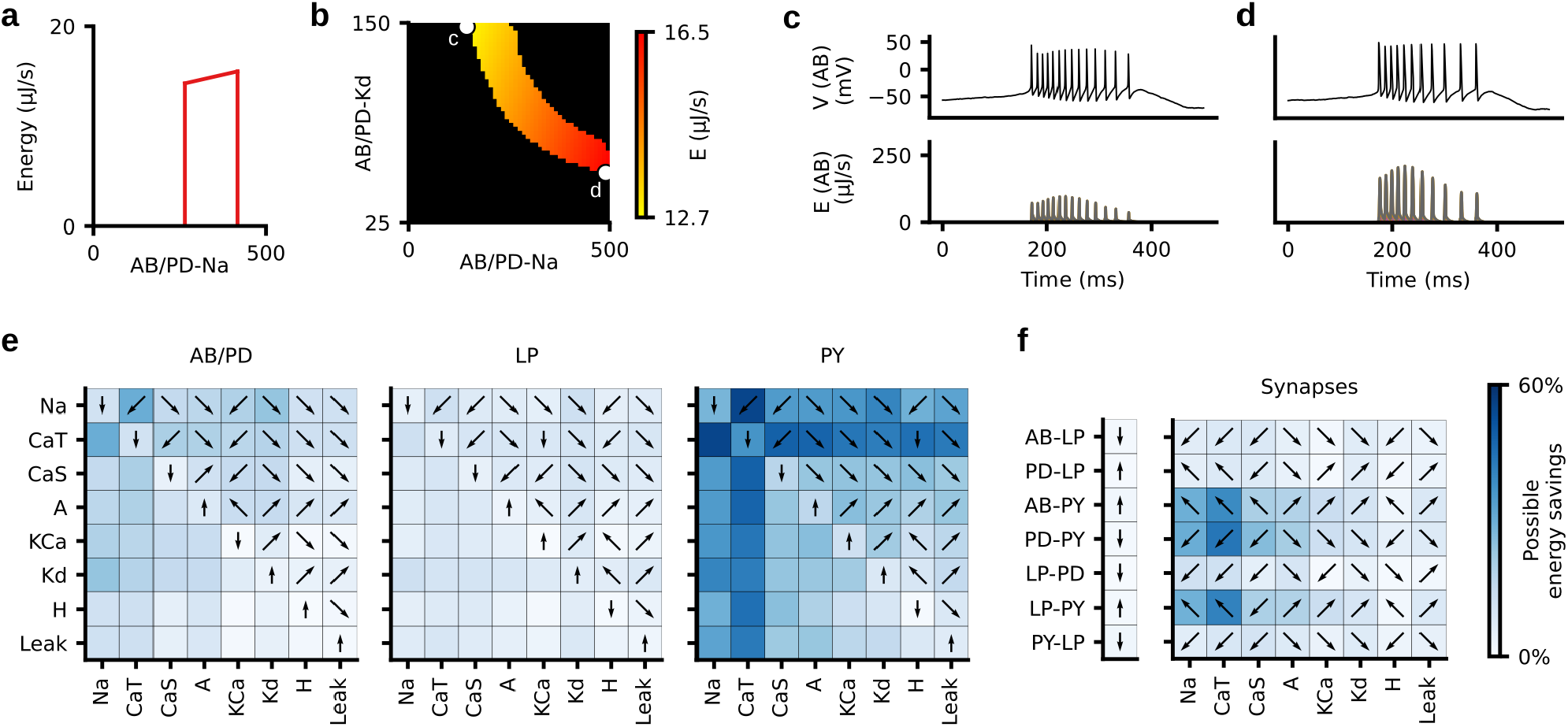
Minimal tuning mechanisms for low energy consumption. (a) Energy consumption (as predicted by linear regression) of several models that differ only in their maximal sodium conductance in the AB/PD neuron. Energy increases with 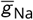. We excluded circuits with too low and too high values of 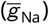, for which the model does not reproduce experimental data. (b) Same as panel (a), but for models that differ in their maximal conductances of sodium (Na) and delayed-rectifier potassium (Kd) in the AB/PD neuron. (c) Top: Voltage trace of the AB/PD neuron for the most efficient configuration within the plane shown in panel (b). Bottom: energy consumption during that activity. (d) Same as panel (c), but for the least efficient configuration. (e) Fraction of energy that can be saved by modifying a single membrane parameter (diagonal of each matrix) or pairs of membrane parameters (upper and lower diagonal). Colorbar as in panel (f). Arrows indicate the direction in which (pairs of) parameters should change in order to reduce energy: Left/right refers to the parameter on the x-axis, top/bottom refers to the parameter on the y-axis. (f) Fraction of energy that can be saved by modifying a synaptic conductance (vector on the left) or the synaptic conductance and one membrane conductance of the respective postsynaptic neuron (matrix on the right).

We repeated this analysis for every conductance and every pair of conductances (Fig. 5e,f). Note that we only considered pairs of parameters within each neuron because pairwise compensation mechanisms across neurons have been shown to be weak in this model [14]. Some of the single-conductance mechanisms can reduce the energy consumption by up to 36%. Pairwise mechanisms, such as reducing the sodium and transient calcium conductances of the PY neuron, can reduce the energy consumption of the entire circuit by up to 55%. When considering only the energy consumed in a specific neuron, pairwise mechanisms can reduce energy consumption by up to 80% (Appendix 1 Fig. 8). Finally, pairwise mechanisms between synapses and conductances of the respective postsynaptic neurons can reduce energy consumption of the entire circuit by up to 43%.

These analyses provide hypotheses for causal mechanisms for how neurons can be tuned into low-energy regimes, while the neural activity keeps satisfying functional constraints. We demonstrated that even simple mechanisms involving one or two conductances can have a substantial impact on the energy consumption of the circuit—thus, low-energy configurations can be found with ‘local’ parameter changes, not requiring fine coordination amongst multiple parameters.

### Neurons can be tuned individually to achieve minimal circuit energy

Next, we asked how single neurons interact to produce functional and efficient circuit activity. Can the energy of the entire circuit be minimized by optimizing the energy of each neuron individually? And does the circuit retain functional activity when neurons are individually optimized for low energy efficiency? Within our database of 35, 939 model configurations, there is a weak correlation between the energies consumed by pairs of neurons, which suggests that the energy consumption between neurons might be independent from one another (Fig. 6a; AB/PD versus LP, correlation coefficient *r* = −0.006, p-value *p* = 0.23; LP versus PY, *r* = 0.02, *p* = 3 · 10^−6^; AB/PD versus PY, *r* = −0.03, *p* = 8 · 10^−9^). We thus investigated whether we could optimize the parameters of each neuron individually for low energy consumption and still retain functional circuit activity. We searched our database of 35,939 model configurations for the single neuron models with minimal energy consumption individually. We selected the five most efficient single neuron parameter combinations for each of the neurons and assembled them into 125 (5^3^) network configurations. We then identified synaptic conductances that match each of these configurations with Markov chain Monte Carlo (Fig. 6b, details in Methods). Notably, given the already estimated full posterior distribution, this step does not require additional simulations.

**Figure 6.**
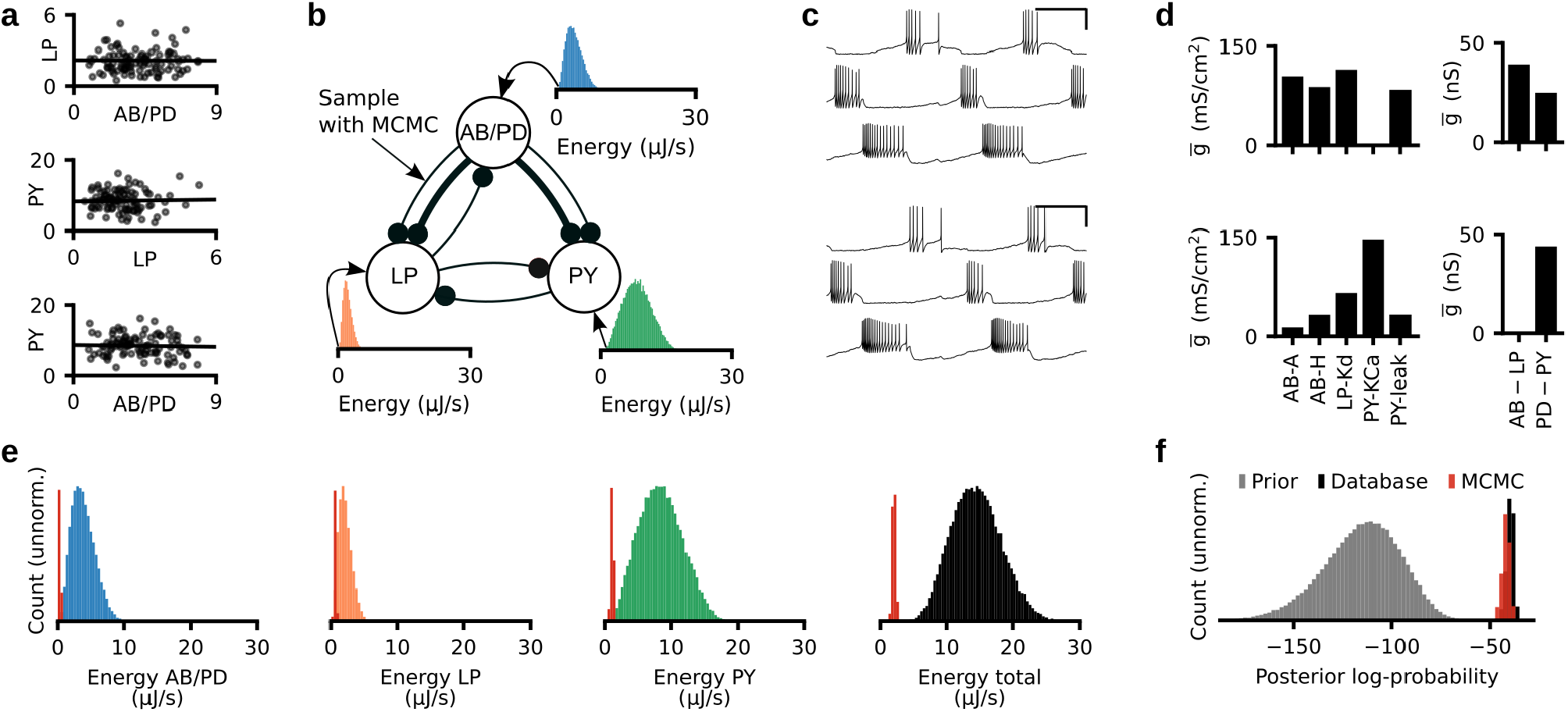
Neurons can be tuned individually to achieve minimal circuit energy consumption. (a) Black dots: Energy consumed by each neuron separately. 100 randomly selected parameter configurations from our database of 35,939 configurations. Black line: Linear regression shows a weak correlation between the energy consumed by pairs of neurons. (b) We select the five most efficient parameter configurations for each neuron separately, and search with Markov chain Monte Carlo (MCMC) for synaptic conductances such that the target circuit activity is achieved. (c) The activity produced by two parameter configurations produced with the strategy described in (b). (d) A subset of the membrane (left) and synaptic (right) conductances for the configurations in (c). Despite generating similar network activity, the configurations have very different circuit parameters. The membrane conductances are scaled with the following factors (left to right): 10,10000,1,100,10000. (e) Histogram over the energy consumption of all 35, 939 models in our database (blue, orange, green, black) and the energy consumption of the configurations produced with the strategy described in panel (b) (red). (f) Histogram of the posterior log-probability for samples from the prior distribution (grey), for the 35,939 models in our database (black), and for the configurations produced with the strategy described in panel (b) (red).

For each of the 125 combinations of membrane conductances, we found a set of synaptic conductances for which the network activity closely resembles experimentally measured activity (Fig. 6c). The resulting configurations have disparate parameters (Fig. 6d) but highly similar network activity. Furthermore, we found that the resulting configurations have similar and very low energy consumption. The energy consumption of these circuits is significantly smaller than that of any of the configurations in our database of 35,939 model configurations (Fig. 6e). This demonstrates that optimizing a specific neuron for energy efficiency does not preclude the connected neurons from being energy efficient. Thus, our results suggest that the pyloric network can be optimized for energy efficiency by tuning neurons individually for low energy consumption.

We estimated how likely are these energy-efficient circuits under the estimated posterior. We found that all these models have similar posterior log-probability as the 35, 939 model configurations in our database (Fig. 6f), i.e. these are as likely to underlie the experimentally measured activity as the database models. Thus, the low-energy configurations were not sampled when generating our original model database because of the high dimensionality of the parameter space, and we cannot exclude the possibility that there might be unsampled regions in parameter space with even more energy-efficient circuit configurations.

### Robustness to temperature does not require an increased metabolic cost

The crab *Cancer borealis* experiences daily and yearly fluctuations in temperature which in turn influence the chemical and physical properties of neurons [32–34]. Nonetheless, neural circuits such as the pyloric network can maintain their functionality in the presence of these temperature variations. As temperature increases, the cycle frequency of the circuit increases exponentially, but the phases between bursts remain relatively constant [35, 39]. We investigated whether the pyloric network trades off robustness to changes in temperature with energy efficiency, i.e. whether temperature-robust solutions are more energetically costly.

The temperature-dependence of a biophysical parameter *R* is captured by the *Q*_10_ value and is defined as follows:

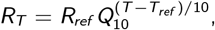

where *R_ref_* is the parameter value at the reference temperature *T_ref_* = 11°C. We extended the model of the pyloric network to include *Q*_10_ values for all maximal membrane and synaptic conductances (details in Methods) [40, 41]. We then used SNPE to identify all maximal membrane and synaptic conductances, as well as the associated *Q*_10_ values (41 parameters in total) that match experimental recordings at 11°C and 27°C (Fig. 7a) [38]. We set the previously identified posterior distribution (Fig. 2e) over circuit parameters given experimental data at 11°C as the new prior distribution, and then applied SNPE to match the model with experimental data at 27°C (Fig. 7b, full posterior in Appendix 1 Fig. 9, details in Methods). We sampled circuit parameters and *Q*_10_ values from the resulting distribution and selected samples whose activity closely matched experimental data at 11°C and 27°C (Fig. 7c). Overall, we generated a database of 967 sets of circuit parameters and *Q*_10_ values. When simulating at temperatures between 11°C and 27°C, these circuits show the characteristic exponential increase in cycle frequency as well as the constant phase relationship between bursts observed experimentally (Fig. 7d) [35].

**Figure 7.**
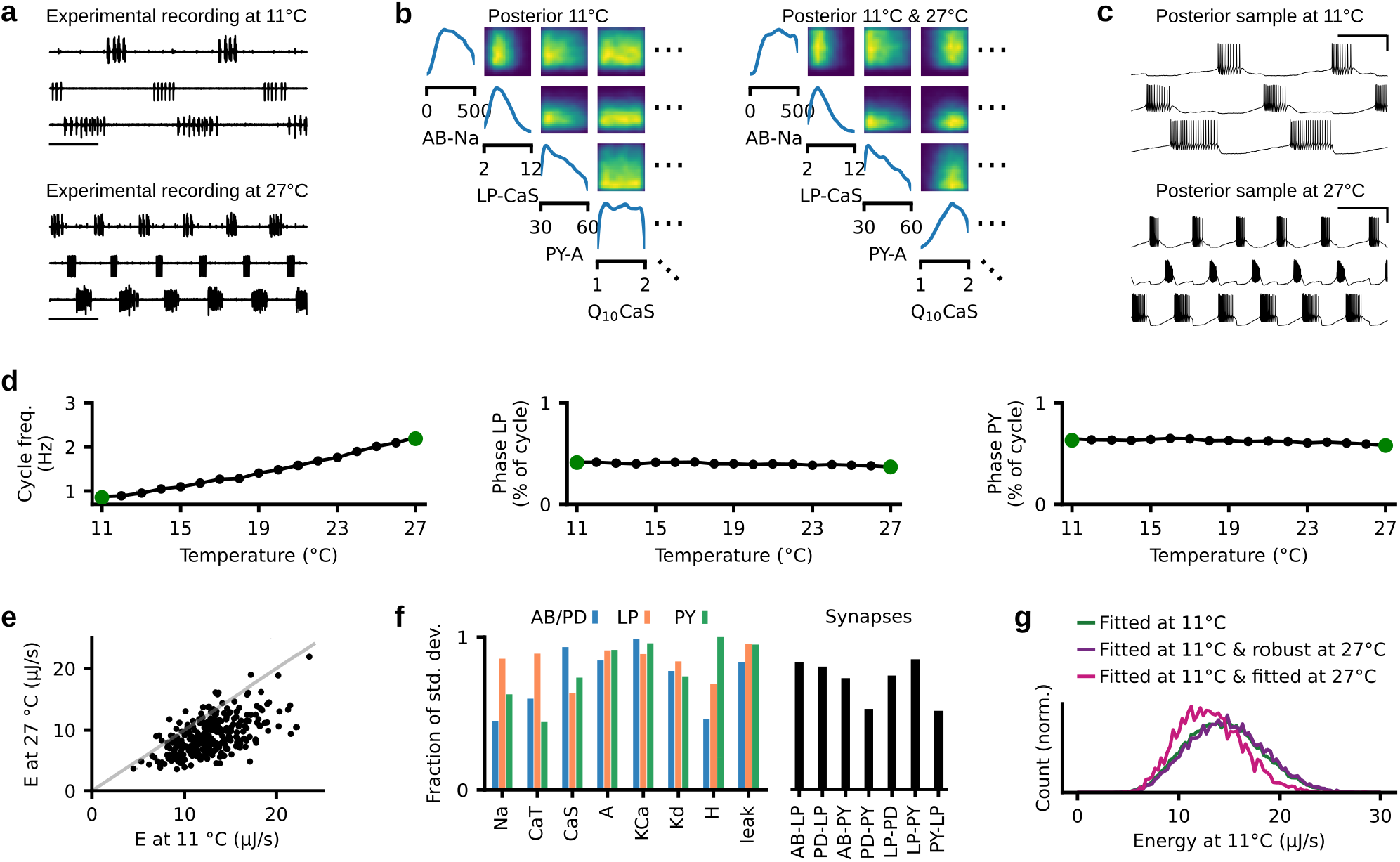
Temperature robustness does not preclude energy efficiency. (a) Top: Experimental data at 11°C. Bottom: Experimental data at 27°C [38]. (b) Left: Posterior distribution given experimental data at 11°C. Right: Posterior given experimental data at 11°C and 27°C. (c) Simulations for a parameter set drawn from the posterior distribution matching experimental data at 11°C and 27°C. Simulations at 11°C (top) and 27°C (bottom). (d) Cycle frequency (left), phase of LP neuron (middle) and phase of PY neuron (right) for parameter set shown in panel (c), simulated at temperatures between 11°C and 27°C. Green dots are the values of the experimental preparations. (e) Energy consumption at 11°C versus 27°C for 967 circuits sampled from the posterior (in (b) right). In grey, the identity line. (f) Standard deviation of parameters for models that match experimental data at 11°C and 27°C and that have energy consumption in the lowest 2% quantile at 11°C and 27°C. Standard deviation is normalized to the standard deviation of the parameters across all 35,939 models in our database. (g) Green: Distribution of the energy consumption of circuits matching experimental data at 11°C. Purple: Distribution of the energy consumption of circuits that match data at 11°C and are robust at 27°C. Pink: Distribution of the energy consumption of circuits that match experimental data at 11°C and 27°C.

We asked whether the energy consumed by the circuit at 11°C is proportional to the energy consumed at 27°C. We found that, despite the number of spikes in our model being higher at higher temperatures, the total energy consumption is lower at 27°C (Fig. 7e; note that, for one of the three preparations, the energy consumptions at 11°C and 27°C are similar; see Appendix 1 Fig. 16). This occurs because at higher temperatures, the increase in the number of spikes is accompanied by an increase in channel time constants and respective decrease in energy per spike (Appendix 1 Fig. 10). In addition, there is a clear correlation between energy consumptions at 11°C and 27°C (Pearson-correlation coefficient: 0.66), although circuit configurations with similar efficiency at 11°C can show a range of energy consumptions at 27°C (Fig. 7e).

We then investigated how the additional constraint of temperature robustness impacts the parameter degeneracy of the pyloric network. We computed the standard deviation of models that match experimental data at 11°C and 27°C and whose energy consumption is in the 2% quantile at both temperatures (Fig. 7f). The resulting standard deviation is smaller than that of all models in our database of 35,939 models, but a large parameter variability remains. Thus, we found a substantial parameter degeneracy in circuits constrained by “pyloric-ness”, energy efficiency and temperature robustness.

Does temperature robustness have an influence on metabolic cost? We computed the energy consumed at 11°C for three different scenarios: First, for all models in our database of 35,939 model configurations matching experimental data recorded at 11°C (same as Fig. 2h). Second, for all models in our database of 35,939 model configurations that are also functional at 27°C (i.e. produce triphasic activity). Third, for all models in our database of 967 model configurations matching experimental data recorded at 11°C and 27°C. In all three of these scenarios, the distribution of metabolic cost was similar (Fig. 7g. Note that the slightly different average energy consumption between the first and the third scenario occurred only in two of the three preparations, see Appendix 1 Fig. 13 and Appendix 1 Fig. 16). In particular, all three scenarios contained configurations that produce energy efficient circuit function. This demonstrates that enforcing temperature robustness does not require the pyloric network to be less energy efficient.

Overall, our analyses indicate that the model of the pyloric network retains substantial parameter degeneracy despite constraints on energy efficiency and temperature robustness. In addition, we showed that temperature robustness does not entail additional metabolic cost.

## Discussion

Neural systems undergo environmental and neuromodulatory perturbations to their mechanisms. The parameter degeneracy of neural systems, i.e. the ability to generate similar activity from disparate parameters, confers a certain degree of robustness to such perturbations [7–10, 42, 43]. However, not all system configurations might be equally desirable, with some configurations being more energy efficient than others [15]. Here, we analysed the energy consumption of parameter configurations with similar activity in the pyloric network of the stomatogastric ganglion. We found that, even when the network activity is narrowly tuned to experimental data, the energy consumption can strongly vary between parameter configurations. Despite this diversity of metabolic costs, energy efficient activity could be produced from a wide range of circuit parameters. When characterising the range of data-consistent parameters, we found a linear relationship between circuit parameters and energy consumption, which allowed us to identify tuning mechanisms for low energy consumption. Lastly, we showed that temperature robustness does not preclude energy efficiency and that parameter degeneracy remains despite metabolic and temperature constraints. These findings were facilitated by a methodological advance that increased the efficiency of previously published tools for simulation-based inference [14, 31, 44, 45].

### Parameter degeneracy under multiple constraints

In addition to a specific activity, neural circuits are likely constrained by other requirements, e.g. low energy consumption or robustness to perturbations such as fluctuations in temperature or pH [35, 40, 41, 46–50]. Here, we investigated how energy efficiency impacts the parameter degeneracy of neural systems. While a plausible hypothesis would have been that energy efficiency reduces or eliminates degeneracy altogether, here we found that parameter degeneracy is preserved, even within circuits with very low energy consumption.

In our work, parameter degeneracy consisted in the range of pyloric-network models that match specific features of experimental activity. We used the same features as in previous work [5], which are physiological constraints of the pyloric network, e.g. cycle duration, burst durations, and gaps and phases of bursts. However, we cannot discard the possibility that the inclusion of additional data-features (e.g. spike height or spike width) would have impacted parameter degeneracy and consequently also the range of energies.

Previous work demonstrated that multiple parameter sets in a model of the AB/PD neuron are temperature robust [40]. Here, we investigated the interplay between energy consumption and temperature robustness at the circuit level, and showed that functional, energy efficient, and temperature robust activity can be generated from disparate circuit parameters. In addition, consistent with previous work in a single neuron model of the grasshopper [28], we found that temperature robustness does not require an increased metabolic cost. Whether these results will generalize with the inclusion of the robustness to additional external perturbations, e.g. pH fluctuations [49, 51], or internal perturbations, e.g. neuromodulation [39], remains a subject for future work.

O’Leary and Marder [52] have demonstrated in a model of the PD neuron that some physiological features (such as duty cycle) can be maintained under temperature perturbations when conductances are scaled by a common factor. We tested the possibility that such invariance under conductance scaling could explain the parameter degeneracy and ranges of energies observed in our circuit model: Scaling the conductances by a common factor would scale the currents and thereby the energy consumption. However, for the parameter ranges we used (similar ranges as in Prinz et al. [5]), scaling the conductances changed physiological features (such as the cycle duration) of the pyloric rhythm and led to the model not fitting the experimental data accurately (Appendix 1 Fig. 4).

More generally, whether there is potential for a system to exhibit parameter degeneracy depends on the number of constraints on the system relative to the number of free parameters: In an over-parameterized system, if there is *any* parameter setting which satisfies the constraints, it is expected that there will be multiple such settings. Our model has 31 conductances and 10 *Q*_10_ values, and we use 18 voltage features at 11°C, one energy consumption constraint and 18 voltage features at 27°. While there is a similar number of constraints relative to the parameter dimensionality, some of those constraints are likely redundant, in which case we have fewer constraints than parameters. Thus, the fact that there are multiple feasible parameter settings is not surprising per se. However, rather than these multiple solutions corresponding to similar parameter values, we found these to be quite disparate in the parameter space.

### Relation to previous work on metabolic cost of neural systems

There has been extensive work on quantifying the metabolic cost of biophysical processes in single neurons [15, 22–26], and how single neurons subject to functional constraints can be tuned to minimize energy consumption [15, 16, 23, 25]. Consistent with this work, we found that total energy consumption of the pyloric network is strongly influenced by the sodium current [25], but also by the transient and slow calcium currents. The maximal sodium conductance is the most prominent driver of the energy per spike: Increases in the conductance lead to an increase of metabolic cost per spike [15,25]. In contrast, calcium currents influence energy consumption through the number of spikes within a burst, despite not consuming much energy themselves. Our results suggest that the maximal conductances of sodium and calcium might be regulated for metabolic efficiency. We thus predict that these conductances are less variable in nature than expected by computational models only matching network activity. Nevertheless, we should note that our findings are based on two simplifying assumptions: First, we studied simple single-compartment neurons rather than more realistic multi-compartment neuron models [53]; and second, the energy measure is derived directly from the Hodgkin-Huxley model [36], rather than taking into account all the complexity of the ionic exchange leading to ATP consumption [15, 21, 23, 25]

Previous studies have demonstrated that synaptic mechanisms can consume a substantial amount of energy [21, 54, 55]. In contrast, in the considered model of the pyloric network, synaptic currents consume only a minor fraction of energy (approximately 0.08% of the total energy is consumed by synapses, whereas Attwell and Laughlin [21] report 40% of energy per action potential being consumed by synaptic mechanisms). This difference is largely due to the low number of connections in the pyloric network [56]: Each neuron projects to up to two other model neurons, whereas the synaptic energy consumption reported in Attwell and Laughlin [21] is based on the assumption of 8000 synaptic boutons per neuron. Thus, models of more complex neural circuits driven by excitatory, recurrent connectivity, such as the ones found in the cortex, might spend a larger fraction of energy on synaptic mechanisms.

### Energy efficiency in the pyloric network

Experimental studies have shown that the parameters of the pyloric network vary across wide ranges [1, 2, 57]. This raises the question of whether these disparate solutions are all tuned for energy efficiency. In our study, we demonstrated that energy-efficient circuit function can be compatible with many parameter configurations. Therefore, despite the variability of the parameters, each configuration in the crab *Cancer Borealis* might be tuned for low energy consumption.

However, the pyloric network is a small subset of the nervous system of the crab and, therefore, likely consumes a small fraction of its total energy budget. Thus, even if the nervous system of the crab is tuned for energy efficiency, it could still achieve this without strict energy requirements for the pyloric network.

### Increasing the efficiency of simulation-based inference

We used a previously introduced tool, SNPE [14, 45] to identify all models consistent with experimentally measured activity as well as prior knowledge about realistic parameter ranges. We improved the efficiency of the method by introducing a classifier that rejects ‘invalid’ simulations [31]. By using this classifier, we were able to improve the accuracy of SNPE while requiring only half as many simulations [14]. Because of this larger simulation-budget, the resulting posterior distributions became more accurate. Furthermore, the trained neural density estimator is amortized, i.e. one can obtain the posterior distribution for multiple experimental preparations without running further simulations or training a new neural network.

The classifier-enhanced SNPE can be applied to other modelling studies in neuroscience. In particular, the classifier to predict ‘invalid’ simulations is valuable whenever there are parameter values for which the computational model of interest produces ill-defined features: E.g. the spike shape cannot be defined in cases where a neuron model does not produce spikes. Our method has the potential to significantly speed up inference in these scenarios.

### Implications for the operation of neural circuits

Our findings suggest that neural circuits can be energy-efficient with largely disparate biophysical parameters, even with highly specific functional requirements under naturally-occurring perturbations. This raises the question of whether such energy efficiency is present in real biological systems, and how these systems could be tuned for metabolic efficiency.

## Acknowledgments

We thank Sara A. Haddad and Eve Marder for sharing their data and discussions, Martin Stemmler for discussions, and Poornima Ramesh, Richard Gao, and Jan Boelts for discussions and comments on the manuscript. The authors thank the International Max Planck Research School for Intelligent Systems (IMPRS-IS) for supporting MD. This work was supported by the German Research Foundation (DFG) through SFB 1089 ‘Synaptic Microcircuits’ and Germany’s Excellence Strategy – EXC-Number 2064/1 – Project number 390727645 as well as the German Federal Ministry of Education and Research (BMBF, project ‘ADIMEM’, FKZ 01IS18052 A-D) to JHM.

## Methods

### Code availability

Code to reproduce the figures is available at https://github.com/mackelab/stg_energy. Code for running SNPE and training a classifier to reject ‘invalid’ simulations is available in our toolbox: https://github.com/mackelab/sbi [58]. A tutorial for how to use these features can be found on our website https://mackelab.org/sbi.

### Data from the crustacean stomatogastric ganglion

We analyzed extracellular recordings of the stomatogastric motor neurons that are involved in the triphasic pyloric rhythm in the crab *Cancer borealis* [38]. The first dataset as seen in Fig. 2 and Fig. 7 is from files 845_082_0044 and 845_082_0064, preparation 1. The second dataset as seen in Appendix 1 Fig. 11 and Appendix 1 Fig. 13 is from files 857_016_0049 and 857_016_0069, preparation 1. The third dataset as seen in Appendix 1 Fig. 14 and Appendix 1 Fig. 16 is from files 845_078_0027 and 845_078_0040, preparation 2. All preparations were decentralized, i.e. the axons of the descending modulatory inputs were severed. The data were recorded at 11°C and 27°C. Full experimental details in Haddad and Marder [39].

### Circuit model of the crustacean stomatogastric ganglion

The circuit model of the crustacean stomatogastric ganglion was adapted from Prinz et al. [5]. The model is composed of three single-compartment neurons, AB/PD, LP, and PY, where the electrically coupled AB and PD neurons are modeled as a single neuron. Each of the model neurons contains 8 currents, a Na^+^ current *I*_Na_, a fast and a slow transient Ca^2+^ current *I*_CaT_ and *I*_CaS_, a transient K^+^ current *I*_A_, a Ca^2+^-dependent K^+^ current *I*_KCa_, a delayed rectifier K^+^ current *I*_Kd_, a hyperpolarization-activated inward current *I*_H_, and a leak current *I*_leak_. In addition, the model contains 7 synapses. As in Prinz et al. [5], these synapses are simulated using a standard model of synaptic dynamics [59]. The synaptic input current into the neurons is given by 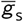, where 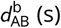. is the maximal synapse conductance, *V*_post_ the membrane potential of the postsynaptic neuron, and *V*_s_ the reversal potential of the synapse. The dynamics of the activation variable *s* are given by

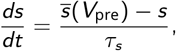

with

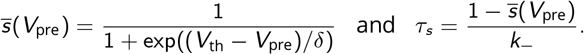

Here, *V*_pre_ is the membrane potential of the presynaptic neuron, *V*_th_ is the half-activation voltage of the synapse, *δ* sets the slope of the activation curve, and *k*_ is the rate constant for transmitter-receptor dissociation rate.

As in Prinz et al. [5], we model two types of synapses, since AB, LP, and PY are glutamatergic neurons whereas PD is cholinergic. We set *E*_s_ = −70 mV and *k*_−_ = 1/40 ms for all glutamatergic synapses and *E*_s_ = −80 mV and *k*_−_ = 1/100 ms for all cholinergic synapses. For both synapse types, we set *V*_th_ = −35 mV and *δ* = 5 mV. The membrane area is 0.628 · 10^−3^ cm^2^.

For each set of membrane and synaptic conductances, we numerically simulate the circuit for 10 seconds with a step size of 0.025 ms. At each time step, each neuron receives Gaussian noise with mean zero and standard deviation 0.001 mV·ms^−05^.

We applied SNPE to infer the posterior over 24 membrane parameters and 7 synaptic parameters, i.e. 31 parameters in total. The 7 synaptic parameters are the maximal conductances of all synapses in the circuit, each of which is varied uniformly in logarithmic domain from 0.01 nS to 1000 nS-with the exception of the synapse from AB to LP, which is varied uniformly in logarithmic domain from 0.01 nS to 10000 nS. The membrane parameters are the maximal membrane conductances for each neuron. The membrane conductances are varied over an extended range of previously reported values [5–14]: The prior distribution over the parameters [Na, CaT, CaS, A, KCa, Kd, H, leak] is uniform with lower bounds *p*_low_ = [0,0, 0,0,0, 25, 0,0] mS cm^−2^ and upper bounds *p*_high_ = [500, 7.5, 8, 60, 15, 150, 0.2, 0.01] mS cm^−2^ for the maximal membrane conductances of the AB neuron, *p*_low_ = [0,0,2,10,0,0,0,0.01] mS cm^−2^ and *p*_high_ = [200, 2.5, 12, 60, 10, 125, 0.06, 0.04] mS cm^−2^ for the maximal membrane conductances of the LP neuron, and *p*_low_ = [0, 0, 0, 30, 0, 50, 0, 0] mS cm^−2^ and *p*_high_ = [600,12.5, 4,60,5,150, 0.06, 0.04] mS cm^−2^ for the maximal membrane conductances of the PY neuron.

We computed 15 summary features proposed by Prinz et al. [5], and 3 additional features [14]. The features proposed by Prinz et al. [5] are 15 salient features of the pyloric rhythm, namely: Cycle period *T* (s), AB/PD burst duration 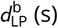, LP burst duration 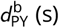, PY burst duration 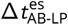, gap AB/PD end to LP start 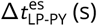 (s), gap LP end to PY start 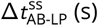, delay AB/PD start to LP start 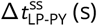, delay LP start to PY start 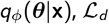, AB/PD duty cycle *d*_AB_, LP duty cycle *d*_LP_, PY duty cycle *d*_PY_, phase gap AB/PD end to LP start Δ_*ϕ*AB-LP_, phase gap LP end to PY start Δ_*ϕ*LP-PY_, LP start phase *ϕ*_LP_, and PY start phase *ϕ*_PY_. Note that several of these values are only defined if each neuron produces rhythmic bursting behavior. In addition, for each of the three neurons, we computed the maximal duration of its voltage being above −30 mV. We did this as we observed—for many model simulations and in contrast with experimental data—long plateaus at around −10 mV during the bursts, and wanted to detect such traces. If the maximal duration was below 5 ms, we set this feature to 5 ms. To extract the summary features from the observed experimental data, we first found spikes by searching for local maxima above a hand-picked voltage threshold, and then extracted the 15 above described features. For the experimental preparation, we set the additional 3 features to 5 ms.

At temperatures higher than 11°C, we include *Q*_10_ values to simulate the biochemical changes of the network parameters. These are defined by an Arrhenius-type factor

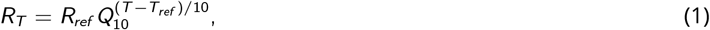

where *R_ref_* is the parameter value at the reference temperature *T_ref_* = 11°C, and *R_T_* is the parameter value at temperature T. Each maximal conductance has a different *Q*_10_, but the *Q*_10_ value is the same across neurons [41]. We introduce one *Q*_10_ for the glutamatergic synapses and one for the cholinergic synapses. The prior distribution for the *Q*_10_ values is a uniform distribution between 1 and 2 for all maximal conductances but the hyperpolarization current, for which the prior bounds are 1 and 4 [35]. The *Q*_10_ values for the time constants are fixed to 2.4 for most *m*-gates and 2.8 for all *h*-gates. Following the results from Caplan et al. [40], the *Q*_10_ values for the m-gates of KCa and CaS as well as for the calcium buffer have lower values: 2.0 for CaS and the calcium buffer and 1.6 for KCa. The *Q*_10_ value for the time constants of the synapses is 1.7.

### Energy consumption

To compute the energy consumption *E* of a specific network activity, we followed the approach of Moujahid et al. [36]. For each neuron, we computed the energy as:

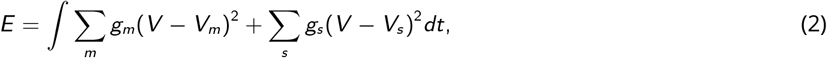

where *g_m_* is the effective conductance of channel *m* (i.e. the product of the respective gating variables, maximal conductance and membrane area) and *g_s_* is the effective synaptic conductance s into the specific neuron. *V_m_* is the reversal potential of the membrane current *m* and *V_s_* is the reversal potential of the synapse *s*. The units of energy are [*S* · *V*^2^ · *s* = *J*], where *S* are Siemens, *V* are Volt, *s* are seconds, and *J* are Joules. The total energy consumption was defined as the sum of the energy consumed by each of the three neurons. Throughout the manuscript, we report the energy per second, which we obtained by dividing the total energy consumption by the duration of the simulation (10 seconds).

The energy per spike was defined as the energy consumed during bursts divided by the respective number of spikes.

### Simulation-based inference

We extended Sequential Neural Posterior Estimation [14] by using a classifier to predict ‘invalid’ simulation outputs. The resulting algorithm is described in algorithm 1.

**Algorithm 1:**
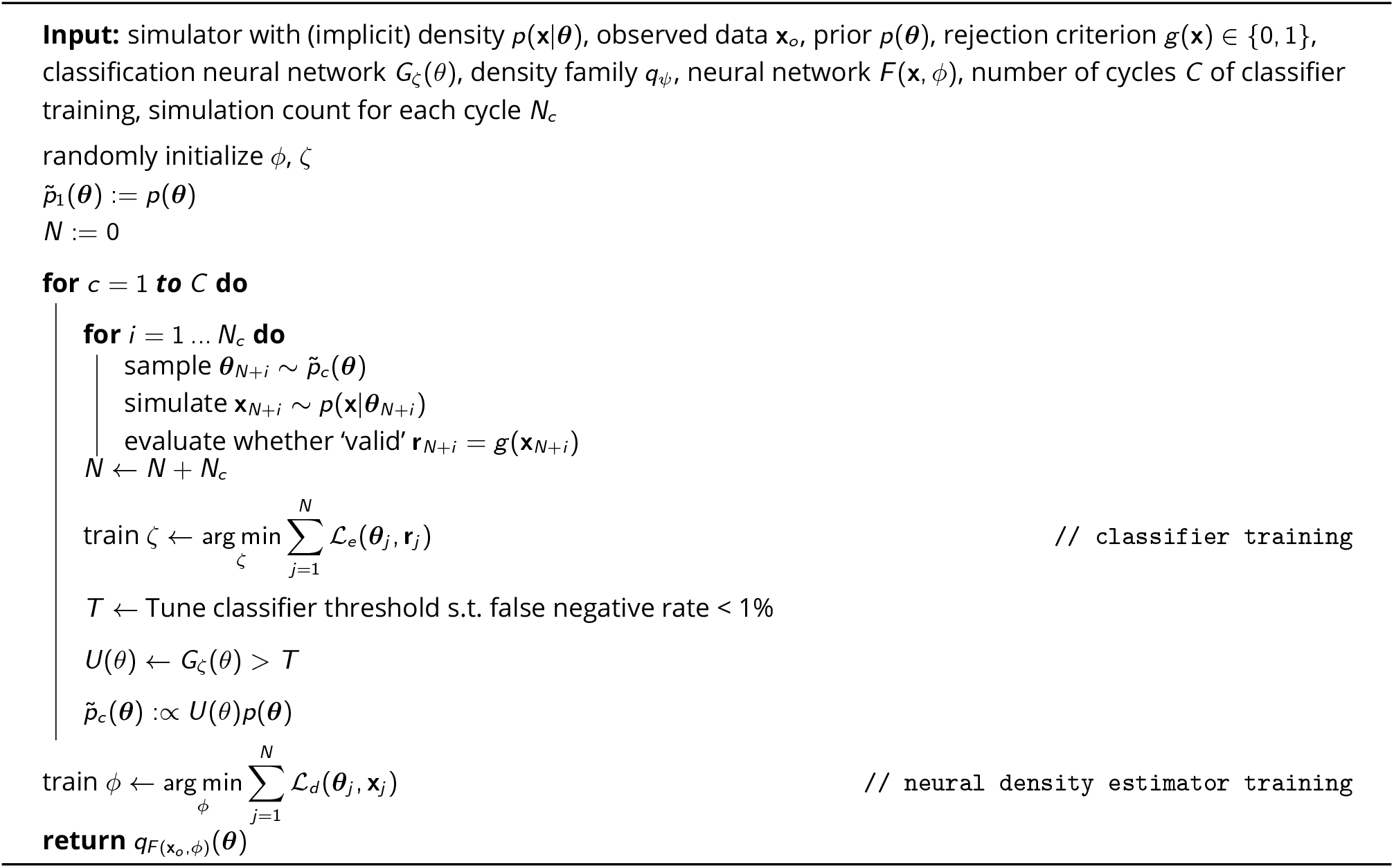
SNPE.

### Proof of convergence of SNPE with classifier

Below, we prove that the posterior distribution inferred by our method converges to the true posterior distribution. SNPE—with the classifier—minimizes the following loss function with respect to the neural network parameters *ϕ*:

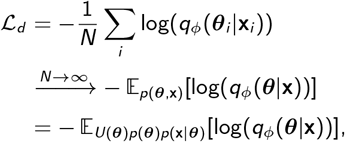

where *U*(***θ***) is a constant *U*(***θ***) = *c* > 0 at least on the posterior support and *U*(***θ***) = 0 elsewhere. Then:

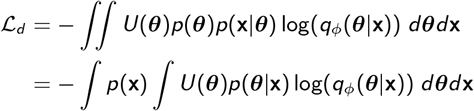

Since *U*(***θ***) > 0 at least on the support of *p*(*θ*|**x**):

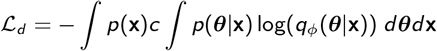

Since the integrand of the integral over ***θ*** is proportional to the Kullback-Leibler-divergence between the true posterior *p*(***θ***|**x**) and the inferred posterior 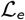 is minimized if and only if *q_ϕ_*(***θ***|**x**) = *p*(***θ***|**x**) for all **x** on the support of *p*(**x**).

### Classifier of ‘valid’ simulations

The algorithm includes a classifier *U*(***θ***) trained to predict ‘valid’ simulations. We use a cross-entropy loss 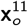. We enforce the classifier *U*(***θ***) to be constant, with *U*(***θ***) = *c* > 0 at least on the posterior support and *U*(***θ***) = 0 elsewhere:

1. In order for *U*(***θ***) to be uniform, we parameterize it as a thresholded binary classifier.
2. To ensure that *c* > 0 at least on the posterior support, we choose the classifier threshold such that there are few false-negatives, i.e. the classifier accepts the parameters if these lie within the posterior support.

If we train the classifier *U*(***θ***) with a large enough number of simulations, so that some are ‘valid’, the trained classifier includes the posterior support. In order to sample from *U*(*θ*)*p*(*θ*), we sample from the prior over parameters *p*(*θ*) and accept the sampled parameters according to the classifier output.

### Inference of the posterior distribution given experimental data at 11°C

Overall, we performed three cycles of simulation and classifier training in order to learn the restricted prior. In the first round, we simulated 3 million parameter sets sampled from the prior. Among these, only 0.97% produced ‘valid’ summary features. We trained a classifier to detect parameter sets leading to ‘valid’ simulation outputs. We used a residual neural network with 80 hidden units, two blocks, a dropout rate of 43%, and a batchsize of 199. To deal with ‘valid’/’invalid’ unbalanced data, we subsampled ‘invalid’ samples in every epoch. We post-hoc tuned the threshold of the classifier such that the ratio of false-negatives was below 1% on a held-out test set. We then drew 3 million samples from the resulting restricted prior. Out of these, 5.17% produced ‘valid’ summary features. We then repeated this procedure and out of 3 million simulations from the resulting restricted prior, 8.45% produced good simulations. Overall, in comparison to Gonçalves et al. [14], we used half as many simulations (9 million versus 18.5 million), but generated a database of ‘valid’ simulations 2.5 times larger. We then used all 438,608 ‘valid’ parameter sets to obtain the posterior distribution with SNPE (see Gonçalves et al. [14] for details). As deep neural density estimator, we chose a neural spline flow(NSF) [60] with 10 transform layers, each consisting of a residual block with two hidden layers, each with 200 hidden units.

Lastly, to ensure that the activity produced by samples from the posterior closely matched experimental data, we sampled 1 million parameter sets from the inferred posterior distribution and performed an additional rejection step, whereby posterior samples had to produce activity within a prescribed distance to the experimental data:

- cycle duration and burst durations deviated from the experimental features by a maximum distance of 0.02 standard deviations of all simulations accepted by the classifier, i.e. 20.6 ms for the cycle duration, and [15.0, 13.5,11.5] ms for the burst durations (of AB/PD, LP, and PY neurons).
- duty cycles, phase gaps, phase delays, and phases deviated from the experimental features by a maximum distance of 0.2 standard deviations.

Out of 1 million samples from the posterior, 35,939 samples fulfilled all these criteria. Notably, these samples are no longer unbiased samples from the posterior distribution as estimated by SNPE, but they make up a database of model configurations whose activity closely matches experimental data.

### Regression neural network

We performed a linear regression to identify the contribution of the circuit parameters to the total energy consumption using scikit-learn [61]. In order to test the robustness of the linear regression findings, we trained a regression network to identify directions in the parameter space predictive of total energy consumption. The regression network had the following characteristics: A Residual Network (ResNet) with one hidden layer with 20 hidden units, ReLU activation functions, and 50% dropout rate [62, 63]. We trained the network with a mean-squared error loss.

After training the regression network, we searched for directions that were most predictive of the network output *f*(·). To do so, we followed the procedure described in Constantine [64] and computed:

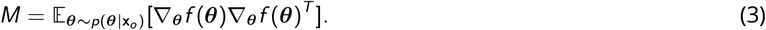

Intuitively, *M* captures how much the regression function *f*(·) changes in different directions of the parameter space, computed as an expected value over posterior samples. We estimated this expected value with a Monte Carlo mean over 10,000 samples from the posterior distribution. We then computed the eigenvalue decomposition of *M*: The eigenvectors of highest eigenvalue are directions in the parameter space along which the output of the regression neural network is most sensitive to changes.

### Minimal tuning mechanisms for low energy consumption

In order to find tuning mechanisms for low energy consumption, we started by varying single or pairs of parameters on a grid while keeping all other parameters constant (at a value taken from our database of 35,939 models). For the matrices in Fig. 5e,f, we repeated the analysis with ten of such models and reported the average.

To predict the energy consumption for any parameter configuration, we used the linear regression model from circuit parameters onto energy consumption described above. To evaluate whether a parameter set matches experimental data, we computed its posterior probability *p*(***θ***|**x**_*o*_). If the posterior probability was above a certain threshold *t*, we considered the parameter set as matching the experimental data, otherwise we discarded the parameter set. To obtain the threshold *t*, we evaluated the posterior probability of all 35,939 models in our database and used the 10% quantile of these probabilities as *t*: within our database of models, 90% of the models would have been considered as matching experimental data, whereas 10% of the models would have been (wrongly) discarded. We evaluated different choices for the quantile used to obtain *t* and obtained qualitatively similar results.

In order to obtain the directions in the parameter space along which energy consumption can be maximally reduced (arrows in Fig. 5e-f), we searched the (1-dimensional or 2-dimensional) grids for the configurations with lowest and highest energy consumption (subject to posterior probability ≥ *t*). For each grid, we reported the sign of the direction between these two points of highest and lowest energy consumption.

### Sampling synaptic conductances, given energy efficient single-neuron configurations

In order to investigate whether efficient single-neuron parameters could lead to efficient and robust network activity, we first searched our database of 35,939 network configurations for the five configurations that had the lowest metabolic cost in each neuron individually. We combined these single neuron configurations to generate 5^3^ = 125 configurations of membrane conductances. For each of the configurations, we then sampled 1,000 synaptic configurations from the distribution:

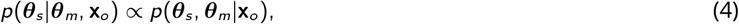

where ***θ***_*s*_ and ***θ***_*m*_ are the synaptic and membrane conductances, respectively. We drew these samples with Markov chain Monte Carlo: Specifically, we used Slice Sampling with axis-aligned updates [65]. We then simulated each of these 5^3^ · 1000 configurations. 72 out of the 5^3^ configurations contained at least one sample that fulfilled our (distance to experimental data) criteria, and 123 configurations contained a sample that fulfilled a slightly wider criteria (allowing twice as much distance from the experimental data). For the remaining two configurations, we drew another 10,000 samples with MCMC and for each of them found at least one configuration whose activity fulfilled the slightly wider criteria. The histograms in figures Fig. 6e-f are produced with all simulations that fulfilled the narrow criteria.

### Posterior distribution given experimental data at 27°C

In order to infer the posterior distribution given experimental data at 27°C, we started by sampling 3 million parameter sets from the 31-dimensional posterior distribution at 11°C:

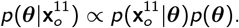

We then drew 3 million sets of *Q*_10_ values from the prior distribution over *Q*_10_ values (*Q*_10_ prior in Methods, Circuit model of the crustacean stomatogastric ganglion). We simulated these 3 million parameter sets at 27°C, from which approximately 18% were ‘valid’ and were used to train a deep neural density estimator (see Proof of convergence of SNPE with classifier). The hyperparameters of the neural density estimator were the same as the ones chosen for the inference at 11°C. Since this density estimator was trained on parameters sampled from the posterior distribution at 11°C, the inferred posterior is an approximation to:

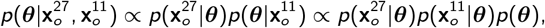

where 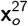 and 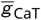 are the features of the experimental data recorded at 11°C and 27°C, respectively. In other words, the resulting posterior distribution matches prior knowledge about circuit parameters as well as experimental data at 11°C and 27°C. Note that we inferred the posterior distribution at 11°C while ignoring the *Q*_10_ values because the *Q*_10_ values, by definition, do not influence the circuit activity at the reference temperature (which is assumed to be 11°C).

### Metabolic efficiency at 27°C

For panels Fig. 7e,f and Fig. 7g (pink plot), we analyzed 967 simulations that closely matched experimental data recorded at 11°C and 27°C. For Fig. 7g (purple plot), we simulated, at 27°C, the 35,939 circuit configurations that match experimental data recorded at 11°C. Out of these, 8121 were robust, i.e. displayed pyloric activity at 27°C.

## Supplementary material

### Supplementary figures

**Supplementary Figure 1.**
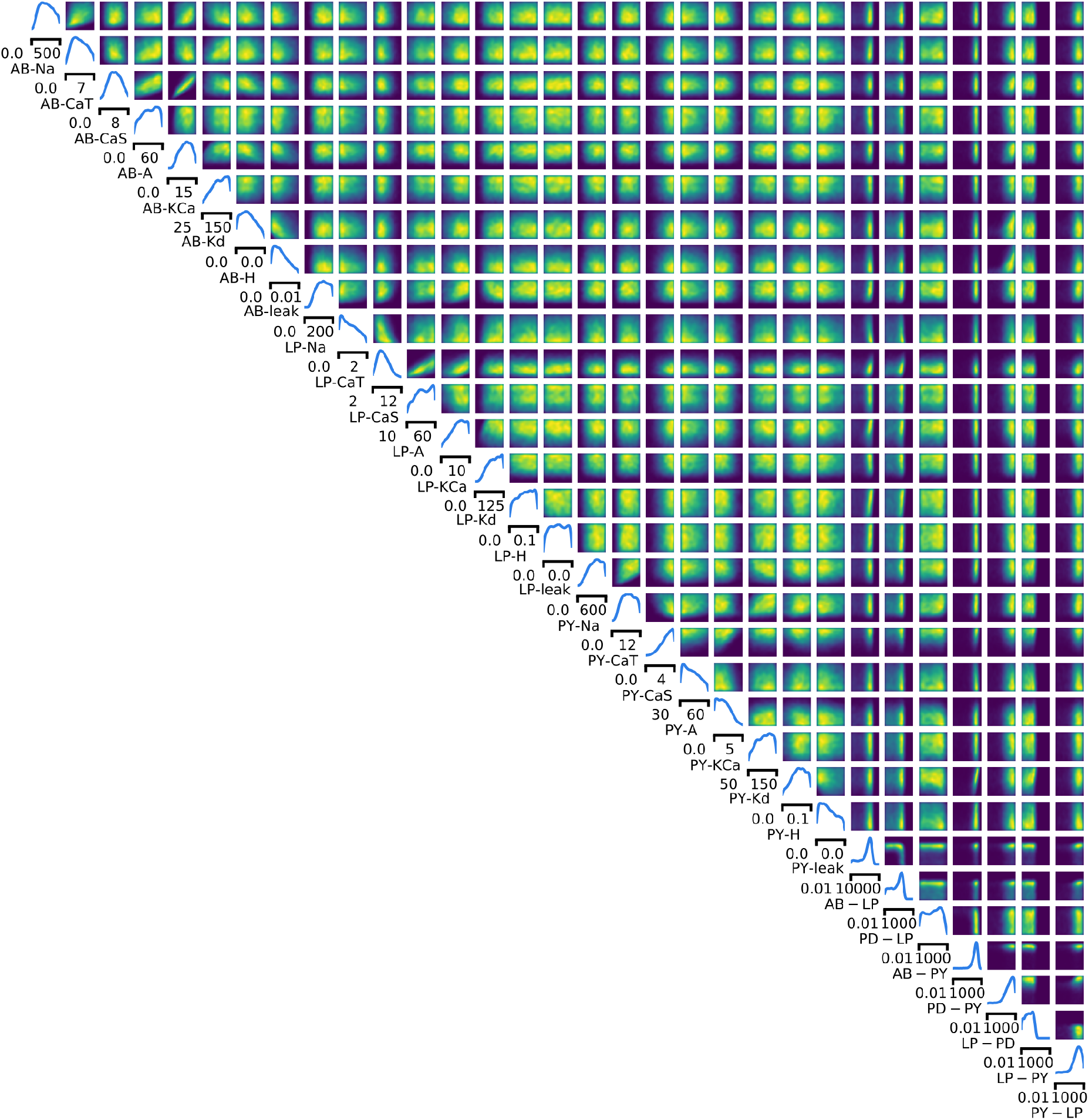
Full posterior distribution over circuit parameters given experimental data at 11°C. Panels on the diagonal are marginals, panels on the upper right are pairwise marginals. The first 24 parameters are membrane conductances, the last 7 parameters are synaptic conductances. All membrane conductances are maximal conductances and are given in mS/cm^2^, all synaptic conductances are given in nS.

**Supplementary Figure 2.**
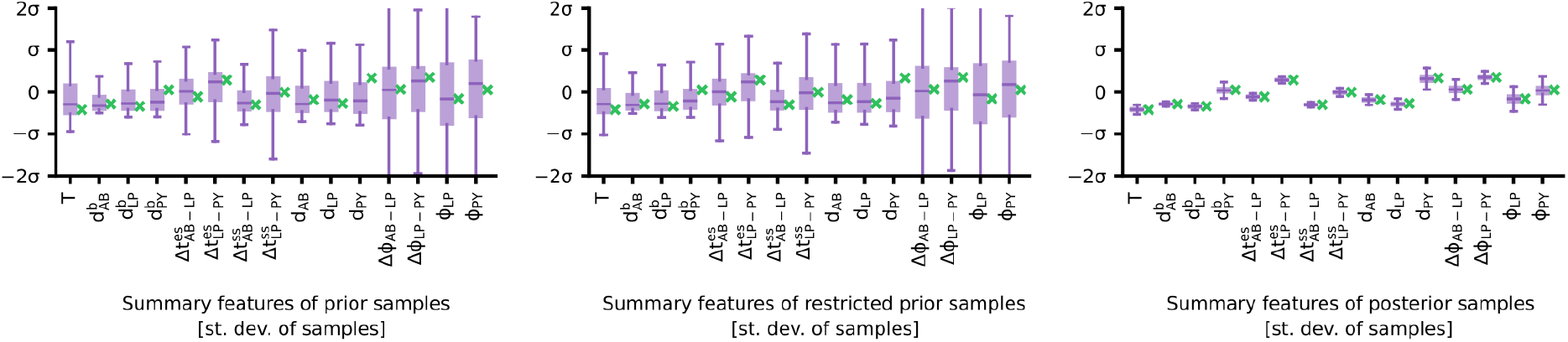
Summary features of activity produced by sampling from the prior, the restricted prior, and the posterior. Experimentally observed activity in green. The boxplots indicate maximum, 75% quantile, median, 25% quantile, and minimum. All summary features are z-scored with the mean and standard deviation of all simulations from prior samples.

**Supplementary Figure 3.**
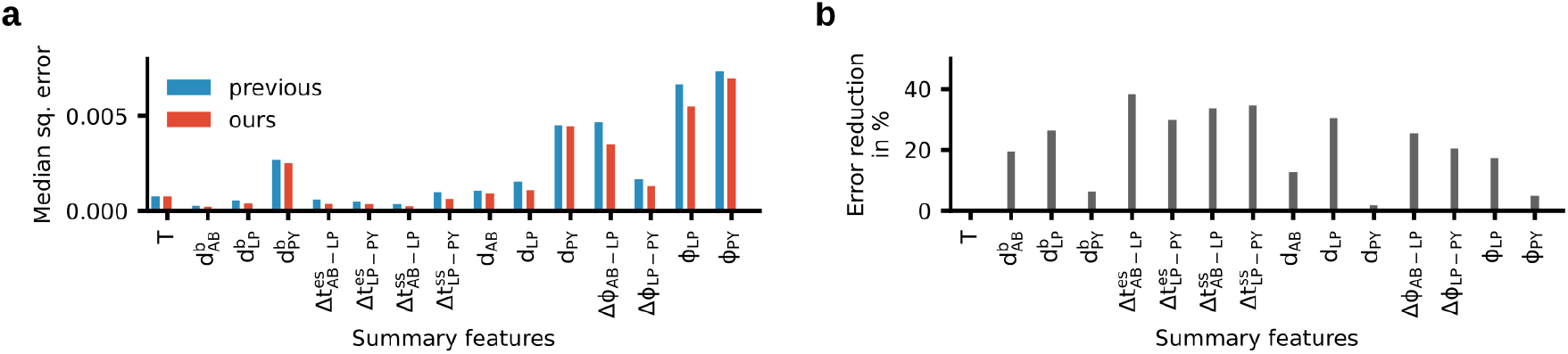
Accuracy of the enhanced version of SNPE versus accuracy in [14]. While we used half as many simulations (9 million versus 18 million), the accuracy of the method improved. (a) Median squared discrepancy between the experimentally measured activity and the activity produced by samples from the posterior. When using the classifier (red), the activity produced by posterior samples is closer to experimental activity than without the classifier (blue). (b) Reduction of mean squared discrepancy between our previous results and the presented method. All distances are computed after z-scoring the summary features with the mean and standard deviation of all prior samples.

**Supplementary Figure 4.**
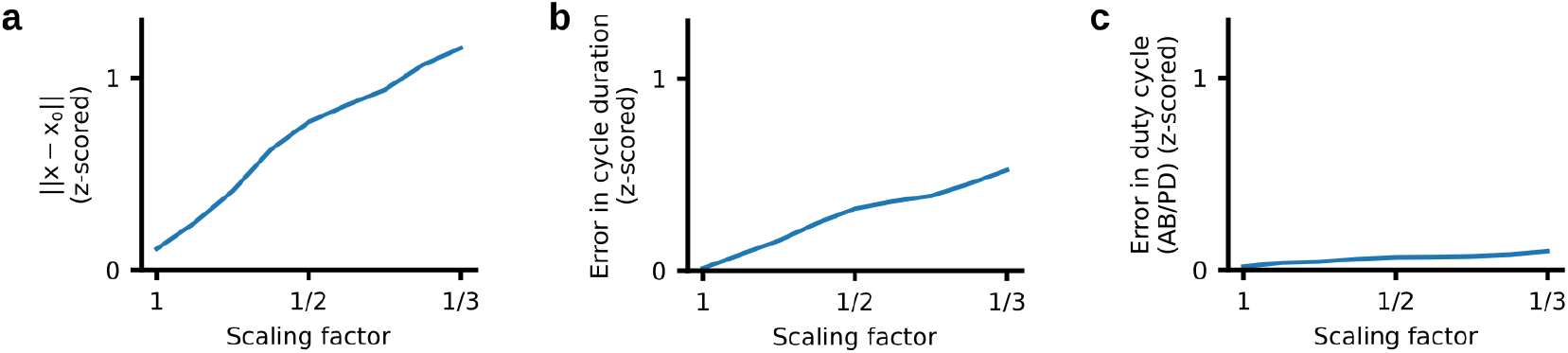
Scaling conductances with a constant factor does not produce activity that matches experimental data. We selected the 20 most expensive circuit configurations from our database of 35,939 models and scaled all conductances by the same factor. The factor ranged from one to 1/3 (x-axis). All errors are z-scored by the standard deviation of prior predictive simulations. (a) As the conductances are scaled down, the average error of the model simulations to the observation increases. (b) The average error between the experimentally observed cycle duration and the simulations increases quickly as the conductances are scaled. (c) The average error between the experimentally observed duty cycle of the AB/PD neuron and the simulations increases minimally as the conductances are scaled.

**Supplementary Figure 5.**
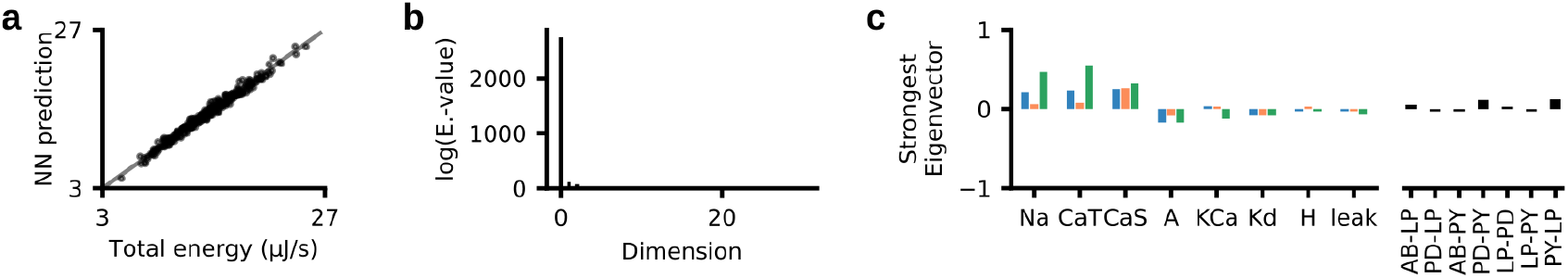
Neural network regression from circuit parameters onto the total energy consumption. (a) Performance of a neural network predicting the total energy from circuit parameters. (b) Eigenvalue-spectrum of the trained neural network reveals a single dominating direction (details in Methods). (c) The eigenvector corresponding to the strongest eigenvalue is similar to the linear regression weights *w* (Fig. 4c).

**Supplementary Figure 6.**
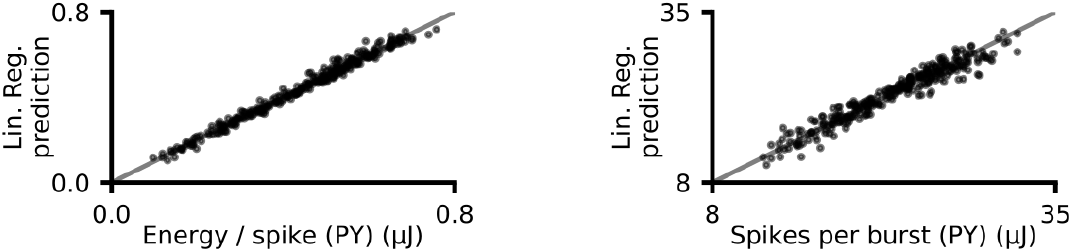
Performance of linear regression. Left: Performance of linear regression from circuit parameters (taken from our database of 35, 393 models) onto energy per spike in the PY neuron. Right: Performance of linear regression from circuit parameters onto the average number of spikes within a burst.

**Supplementary Figure 7.**
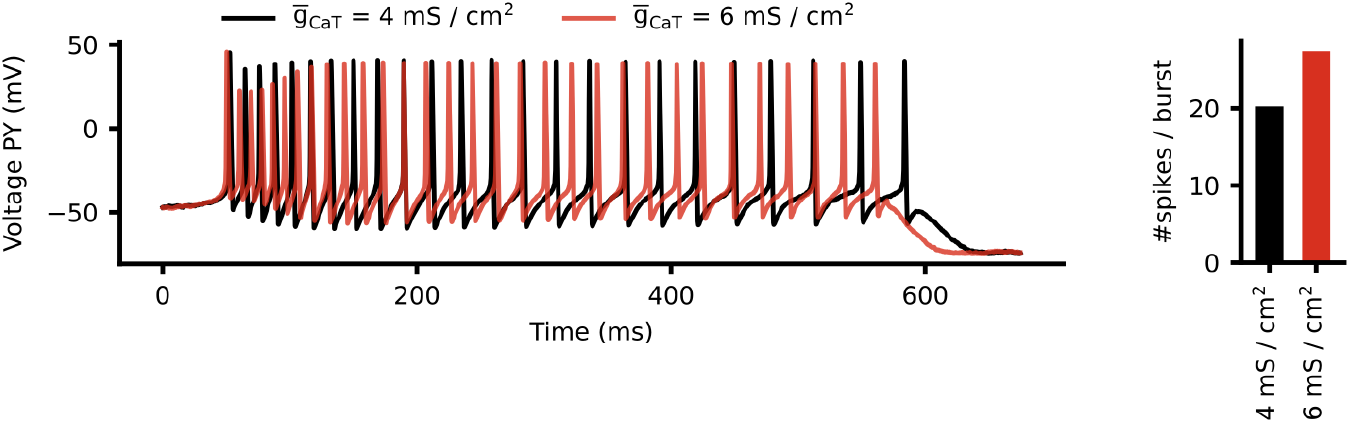
Influence of maximal conductance of transient calcium current on circuit function. Left: Voltage trace in the PY neuron during activity produced by two circuit configurations (black and red) which are identical apart from the magnitude of 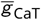. Right: The average number of spikes per burst in the PY neuron for the two configurations. The configuration with higher 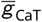 produces more spikes.

**Supplementary Figure 8.**
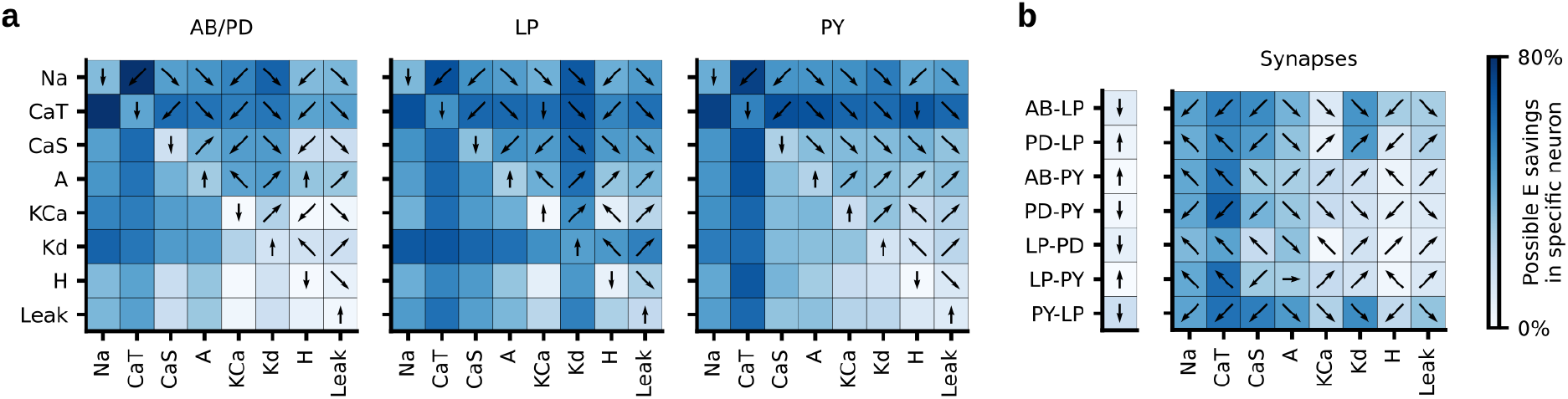
Potential energy savings in a single neuron. (a) The fraction of energy that can be saved in a single neuron by modifying just a single membrane parameter (diagonal of each matrix) or pairs of membrane parameters (upper and lower diagonal) in that neuron. Unlike Fig. 5e, only the energy in the neuron whose parameters are varied is considered. Colorbar as in panel (b). The arrows indicate the direction in which (pairs of) parameters should change in order to reduce energy: Left/right refers to the parameter on the x-axis, top/bottom refers to the parameter on the y-axis. (b) The fraction of energy that can be saved by modifying a synaptic conductance (vector on the left) or the synaptic conductance and one membrane conductance of the postsynaptic neuron (matrix on the right). The energy is computed in the postsynaptic neuron.

**Supplementary Figure 9.**
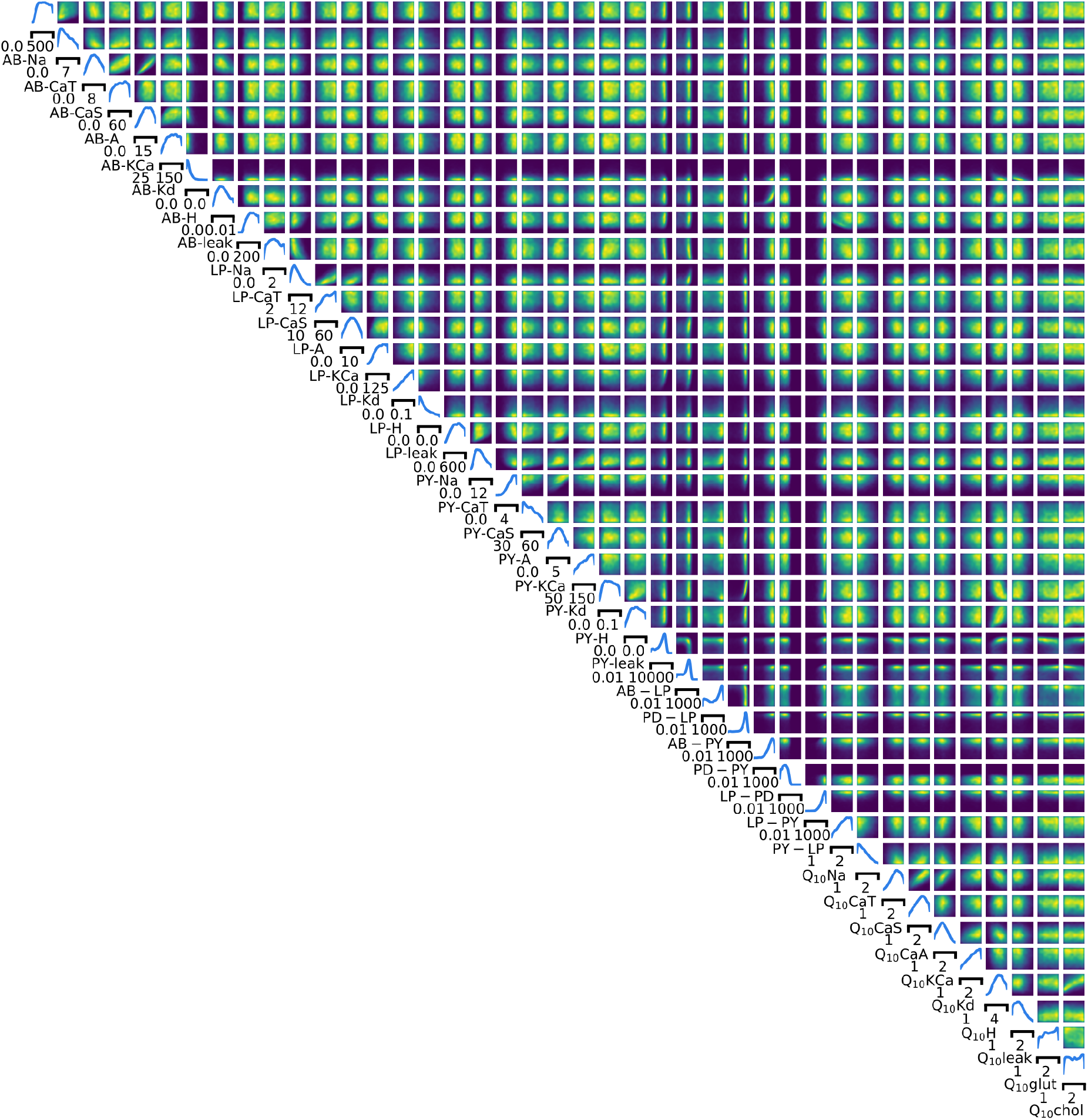
**Full posterior distribution over 31 circuit parameters and 10** *Q*_10_ **parameters given experimental data at 11°C and 27°C.**

**Supplementary Figure 10.**
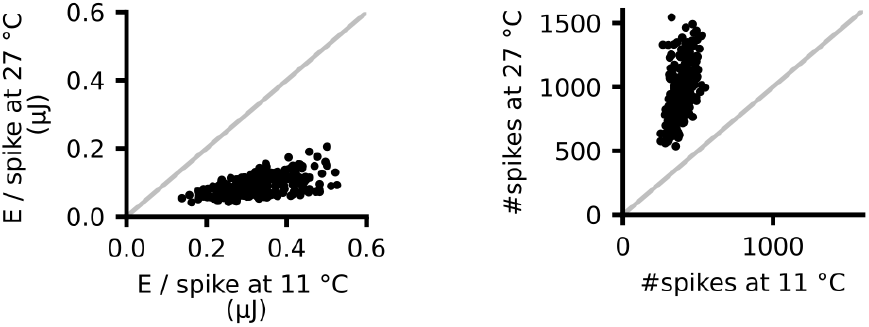
Energy consumption of the circuit at 11°C and at 27°C. Energy per spike (left) and number of spikes (right) for parameter configurations simulated at 11°C and 27°C. The energy per spike is smaller at higher temperatures, but the number of spikes is higher at higher temperatures.

**Supplementary Figure 11.**
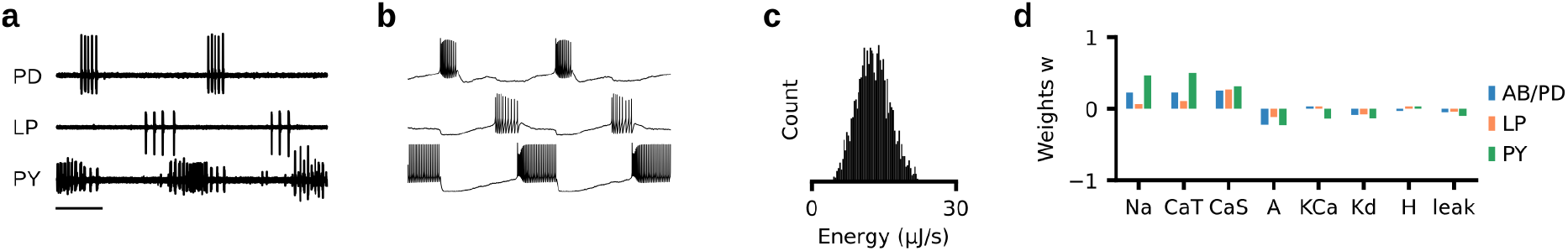
Analysis of a second experimental preparation. (a) Experimental data recorded at 11°C. (b) Sample from posterior distribution matches the experimental data. (c) Energy consumption of 2804 model configurations that closely match experimental data. (d) Weights *w* of a linear regression from circuit parameters onto total energy consumption.

**Supplementary Figure 12.**
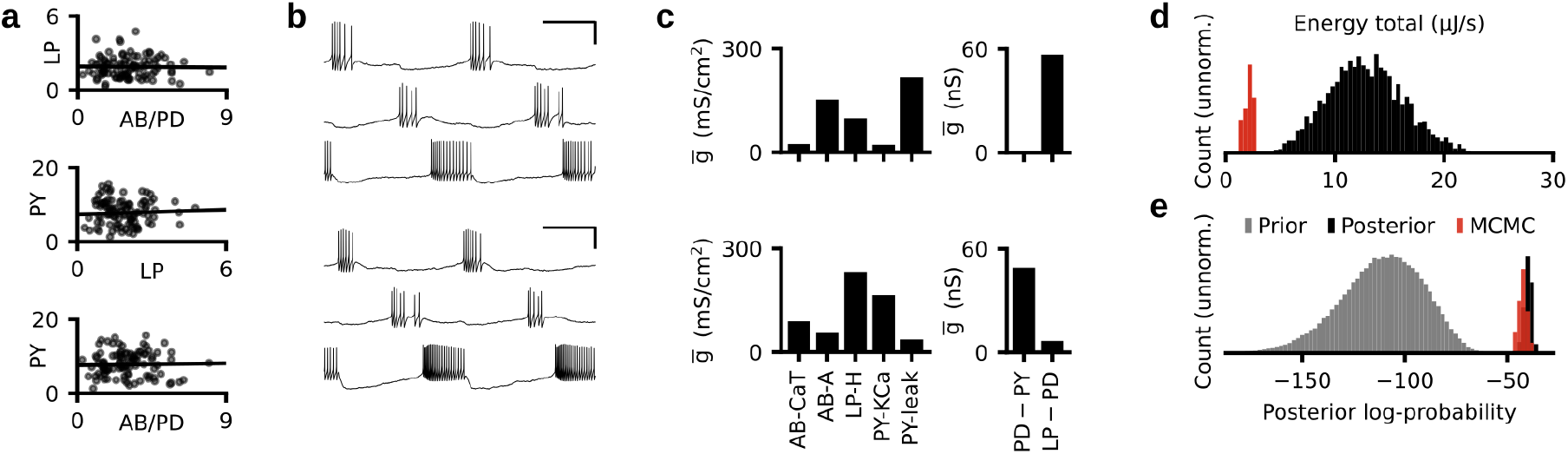
Tuning neurons individually, for a second experimental preparation. (a) Black dots: Energy consumed by each neuron separately. Black line: Linear regression (correlation coefficient *r* = −0.009, p-value *p* = 0.39; LP versus PY, *r* = 0.21, *p* = 0.0012; AB/PD versus PY, *r* = 0.059, *p* = 0.10). (b) The activity produced by two parameter configurations produced with the strategy described in Fig. 6b. (c) A subset of the membrane (left) and synaptic (right) conductances for the configurations in panel (c). The membrane conductances are scaled with the following factors (left to right): 100,10,10,000,100,10,000. (d) Histogram over the total energy consumption of all 2804 model configurations in our database and the energy consumption of the configurations produced with the strategy described in Fig. 6b (red). (e) Histogram of the posterior log-probability for samples from the prior distribution (grey), for the 2804 models in our database (black), and for the configurations produced with the strategy described in Fig. 6b (red).

**Supplementary Figure 13.**
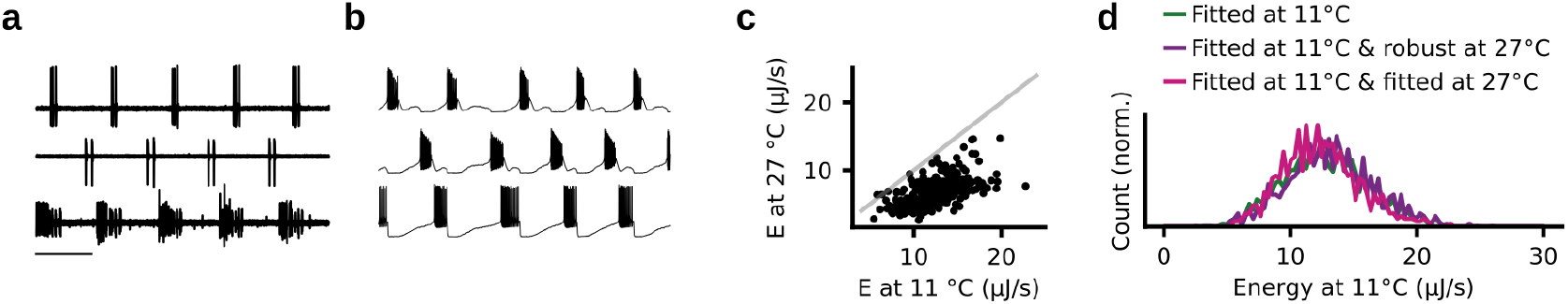
Analysis of temperature robustness of a second experimental preparation. (a) Experimental data recorded at 27°C. (b) Sample from posterior distribution matches the experimental data. (c) Energy consumption at 11°C versus energy consumption at 27°C. (d) Green: Distribution of the energy consumption of circuits matching experimental data at 11°C. Purple: Distribution of the energy consumption of circuits that match data at 11°C and are robust at 27°C. Pink: Distribution of the energy consumption of circuits that match experimental data at 11°C and 27°C.

**Supplementary Figure 14.**
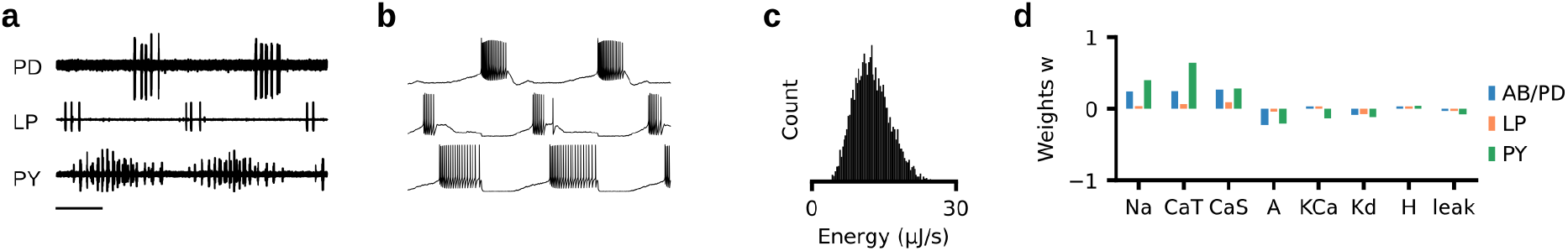
Analysis of a third experimental preparation. (a) Experimental data recorded at 11°C. (b) Sample from posterior distribution matches the experimental data. (c) Energy consumption of 6926 model configurations that closely match experimental data. (d) Weights *w* of a linear regression from circuit parameters onto total energy consumption.

**Supplementary Figure 15.**
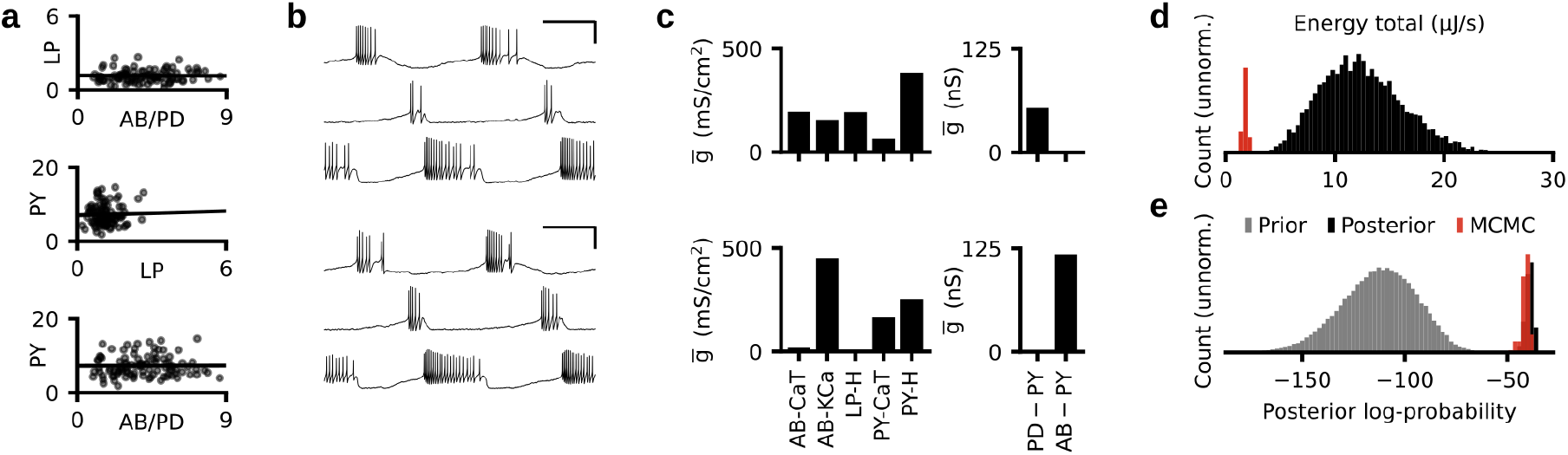
Tuning neurons individually, for a third experimental preparation. (a) Black dots: Energy consumed by each neuron separately. Black line: Linear regression (correlation coefficient *r* = −0.003, p-value *p* = 0.39; LP versus PY, *r* = 0.19, *p* = 0.007; AB/PD versus PY, *r* = 0.006, *p* = 0.77). (b) The activity produced by two parameter configurations produced with the strategy described in Fig. 6b. (c) A subset of the membrane (left) and synaptic (right) conductances for the configurations in panel (c). The membrane conductances are scaled with the following factors (left to right): 100,100,10,000,100,10,000. (d) Histogram over the total energy consumption of all 6926 model configurations in our database and the energy consumption of the configurations produced with the strategy described in Fig. 6b (red). (e) Histogram of the posterior log-probability for samples from the prior distribution (grey), for the 6926 models in our database (black), and for the configurations produced with the strategy described in Fig. 6b (red).

**Supplementary Figure 16.**
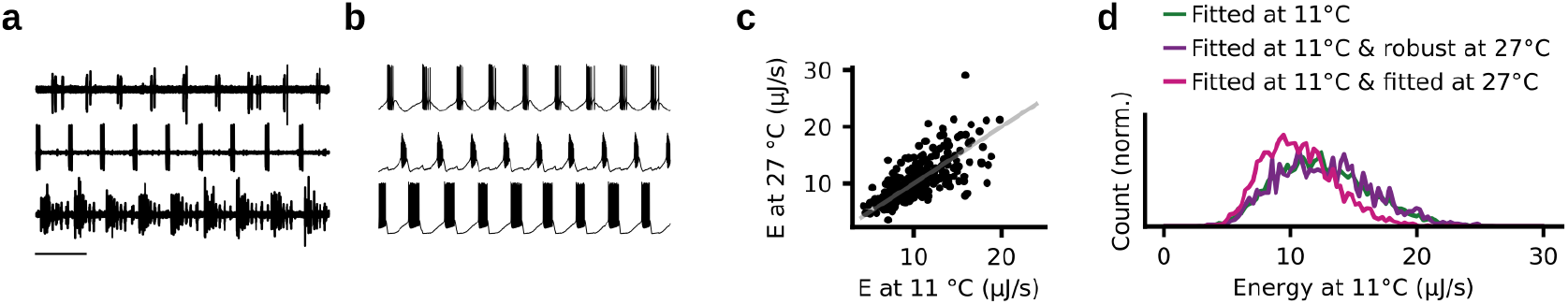
Analysis of temperature robustness of a third experimental preparation. (a) Experimental data recorded at 27°C. (b) Sample from posterior distribution matches the experimental data. (c) Energy consumption at 11°C versus energy consumption at 27°C. (d) Green: Distribution of the energy consumption of circuits matching experimental data at 11°C. Purple: Distribution of the energy consumption of circuits that match data at 11°C and are robust at 27°C. Pink: Distribution of the energy consumption of circuits that match experimental data at 11°C and 27°C.

## References

[1] J. Golowasch, L. Abbott, and E. Marder. Activity-dependent regulation of potassium currents in an identified neuron of the stomatogastric ganglion of the crab cancer borealis. Journal of Neuroscience, 19(20):RC33–RC33, 1999.

[2] J. Golowasch, M. S. Goldman, L. Abbott, and E. Marder. Failure of averaging in the construction of a conductance-based neuron model. Journal of neurophysiology, 87(2):1129–1131, 2002.

[3] M. S. Goldman, J. Golowasch, E. Marder, and L. Abbott. Global structure, robustness, and modulation of neuronal models. Journal of Neuroscience, 21(14):5229–5238, 2001.

[4] A. A. Prinz, C. P. Billimoria, and E. Marder. Alternative to hand-tuning conductance-based models: construction and analysis of databases of model neurons. Journal of Neurophysiology, 90(6):3998–4015, 2003.

[5] A. A. Prinz, D. Bucher, and E. Marder. Similar network activity from disparate circuit parameters. Nature Neuroscience, 7(12): 1345, 2004.

[6] R. C. Roffman, B. J. Norris, and R. L. Calabrese. Animal-to-animal variability of connection strength in the leech heartbeat central pattern generator. Journal of neurophysiology, 107(6):1681–1693, 2012.

[7] G. M. Edelman and J. A. Gally. Degeneracy and complexity in biological systems. Proceedings of the National Academy of Sciences, 98(24):13763–13768, 2001.

[8] E. Marder and A. L. Taylor. Multiple models to capture the variability in biological neurons and networks. Nature Neuroscience, 14(2):133, 2011.

[9] J. N. MacLean, Y. Zhang, B. R. Johnson, and R. M. Harris-Warrick. Activity-independent homeostasis in rhythmically active neurons. Neuron, 37(1):109–120, 2003.

[10] J. N. MacLean, Y. Zhang, M. L. Goeritz, R. Casey, R. Oliva, J. Guckenheimer, and R. M. Harris-Warrick. Activity-independent coregulation of ia and ih in rhythmically active neurons. Journal of Neurophysiology, 94(5):3601–3617, 2005.

[11] R. N. Gutenkunst, J. J. Waterfall, F. P. Casey, K. S. Brown, C. R. Myers, and J. P. Sethna. Universally sloppy parameter sensitivities in systems biology models. PLoS Computational Biology, 3(10):e189, 2007.

[12] R. Grashow, T. Brookings, and E. Marder. Compensation for variable intrinsic neuronal excitability by circuit-synaptic interactions. Journal of Neuroscience, 30(27):9145–9156, 2010.

[13] T. O’Leary, A. C. Sutton, and E. Marder. Computational models in the age of large datasets. Current Opinion in Neurobiology, 32: 87–94, 2015.

[14] P. J. Gonçalves, J.-M. Lueckmann, M. Deistler, M. Nonnenmacher, K. Öcal, G. Bassetto, C. Chintaluri, W. F. Podlaski, S. A. Haddad, T. P. Vogels, et al. Training deep neural density estimators to identify mechanistic models of neural dynamics. Elife, 9:e56261, 2020.

[15] A. Hasenstaub, S. Otte, E. Callaway, and T. J. Sejnowski. Metabolic cost as a unifying principle governing neuronal biophysics. Proceedings of the National Academy of Sciences, 107(27):12329–12334, 2010.

[16] B. Sengupta, A. A. Faisal, S. B. Laughlin, and J. E. Niven. The effect of cell size and channel density on neuronal information encoding and energy efficiency. Journal of Cerebral Blood Flow &amp; Metabolism, 33(9):1465–1473, 2013.

[17] B. Sengupta and M. B. Stemmler. Power consumption during neuronal computation. Proceedings of the IEEE, 102(5):738–750, 2014.

[18] J. Astrup, P. M. Sørensen, and H. R. Sørensen. Oxygen and glucose consumption related to na+-k+ transport in canine brain. Stroke, 12(6):726–730, 1981.

[19] J. Astrup, P. M. Sørensen, and H. R. Sørensen. Inhibition of cerebral oxygen and glucose consumption in the dog by hypothermia, pentobarbital, and lidocaine. Anesthesiology: The Journal of the American Society of Anesthesiologists, 55(3):263–268, 1981.

[20] L. Sokoloff. Energetics of functional activation in neural tissues. Neurochemical research, 24(2):321–329, 1999.

[21] D. Attwell and S. B. Laughlin. An energy budget for signaling in the grey matter of the brain. Journal of Cerebral Blood Flow & Metabolism, 21(10):1133–1145, 2001.

[22] H. Alle, A. Roth, and J. R. Geiger. Energy-efficient action potentials in hippocampal mossy fibers. Science, 325(5946):1405–1408, 2009.

[23] M. B. Stemmler, B. Sengupta, S. Laughlin, and J. Niven. Energetically optimal action potentials. In Advances in neural information processing systems, pages 1566–1574, 2011.

[24] B. C. Carter and B. P. Bean. Sodium entry during action potentials of mammalian neurons: incomplete inactivation and reduced metabolic efficiency in fast-spiking neurons. Neuron, 64(6):898–909, 2009.

[25] B. Sengupta, M. Stemmler, S. B. Laughlin, and J. E. Niven. Action potential energy efficiency varies among neuron types in vertebrates and invertebrates. PLoS Comput Biol, 6(7):e1000840, 2010.

[26] G. Yi, Y. Fan, and J. Wang. Metabolic cost of dendritic ca2+ action potentials in layer 5 pyramidal neurons. Frontiers in neuroscience, 13, 2019.

[27] S. Onasch and J. Gjorgjieva. Circuit stability to perturbations reveals hidden variability in the balance of intrinsic and synaptic conductances. Journal of Neuroscience, 40(16):3186–3202, 2020.

[28] F. A. Roemschied, M. J. Eberhard, J.-H. Schleimer, B. Ronacher, and S. Schreiber. Cell-intrinsic mechanisms of temperature compensation in a grasshopper sensory receptor neuron. Elife, 3:e02078, 2014.

[29] R. M. Harris-Warrick, E. Marder, A. I. Selverston, M. Moulins, T. J. Sejnowski, and T. A. Poggio. Dynamic biological networks: the stomatogastric nervous system. MIT press, 1992.

[30] E. Marder and D. Bucher. Understanding circuit dynamics using the stomatogastric nervous system of lobsters and crabs. Annu. Rev. Physiol., 69:291–316, 2007.

[31] J.-M. Lueckmann, P. J. Goncalves, G. Bassetto, K. Öcal, M. Nonnenmacher, and J. H. Macke. Flexible statistical inference for mechanistic models of neural dynamics. In Advances in Neural Information Processing Systems, pages 1289–1299, 2017.

[32] L. Stehlik, C. MacKenzie Jr, and W. Morse. Distribution and abundance of four brachyuran crabs on the northwest atlantic shelf. Fishery Bulletin, 89(3):473–492, 1991.

[33] M. J. Donahue, A. Nichols, C. A. Santamaria, P. E. League-Pike, C. J. Krediet, K. O. Perez, and M. J. Shulman. Predation risk, prey abundance, and the vertical distribution of three brachyuran crabs on gulf of maine shores. Journal of Crustacean Biology, 29(4):523–531, 2009.

[34] C. J. Krediet and M. J. Donahue. Growth-mortality trade-offs along a depth gradient in cancer borealis. Journal of Experimental Marine Biology and Ecology, 373(2):133–139, 2009.

[35] L. S. Tang, M. L. Goeritz, J. S. Caplan, A. L. Taylor, M. Fisek, and E. Marder. Precise temperature compensation of phase in a rhythmic motor pattern. PLoS Biol, 8(8):e1000469, 2010.

[36] A. Moujahid, A. d’Anjou, F. Torrealdea, and F. Torrealdea. Energy and information in hodgkin-huxley neurons. Physical Review E, 83(3):031912, 2011.

[37] L. M. Alonso and E. Marder. Visualization of currents in neural models with similar behavior and different conductance densities. eLife, 8:e42722, 2019.

[38] S. A. Haddad and E. Marder. Recordings from the c. borealis stomatogastric nervous system at different temperatures in the decentralized condition, July 2021. URL https://doi.org/10.5281/zenodo.5139650.

[39] S. A. Haddad and E. Marder. Circuit robustness to temperature perturbation is altered by neuromodulators. Neuron, 100(3): 609–623, 2018.

[40] J. S. Caplan, A. H. Williams, and E. Marder. Many parameter sets in a multicompartment model oscillator are robust to temperature perturbations. Journal of Neuroscience, 34(14):4963–4975, 2014.

[41] L. M. Alonso and E. Marder. Temperature compensation in a small rhythmic circuit. Elife, 9:e55470, 2020.

[42] E. Marder. Variability, compensation, and modulation in neurons and circuits. Proceedings of the National Academy of Sciences, 108(Supplement 3):15542–15548, 2011.

[43] E. Marder, M. L. Goeritz, and A. G. Otopalik. Robust circuit rhythms in small circuits arise from variable circuit components and mechanisms. Current Opinion in Neurobiology, 31:156–163, 2015.

[44] G. Papamakarios, T. Pavlakou, and I. Murray. Masked autoregressive flow for density estimation. In Advances in Neural Information Processing Systems, pages 2338–2347, 2017.

[45] D. Greenberg, M. Nonnenmacher, and J. Macke. Automatic posterior transformation for likelihood-free inference. In International Conference on Machine Learning, pages 2404–2414, 2019.

[46] L. S. Tang, A. L. Taylor, A. Rinberg, and E. Marder. Robustness of a rhythmic circuit to short-and long-term temperature changes. Journal of Neuroscience, 32(29):10075–10085, 2012.

[47] A. Rinberg, A. L. Taylor, and E. Marder. The effects of temperature on the stability of a neuronal oscillator. PLoS Comput Biol, 9 (1):e1002857, 2013.

[48] T. O’Leary and E. Marder. Temperature-robust neural function from activity-dependent ion channel regulation. Current Biology, 26(21):2935–2941, 2016.

[49] J. A. Haley, D. Hampton, and E. Marder. Two central pattern generators from the crab, cancer borealis, respond robustly and differentially to extreme extracellular ph. Elife, 7:e41877, 2018.

[50] S. Gorur-Shandilya, E. M. Cronin, A. C. Schneider, S. A. Haddad, P. Rosenbaum, D. Bucher, F. Nadim, and E. Marder. Mapping circuit dynamics during function and dysfunction. bioRxiv, 2021.

[51] J. Ratliff, A. Franci, E. Marder, and T. O’Leary. Neuronal oscillator robustness to multiple global perturbations. Biophysical Journal, 2021.

[52] T. O’Leary and E. Marder. Temperature-robust neural function from activity-dependent ion channel regulation. Current Biology, 26(21):2935–2941, 2016.

[53] G. Le Masson, S. Przedborski, and L. Abbott. A computational model of motor neuron degeneration. Neuron, 83(4):975–988, 2014.

[54] W. B. Levy and R. A. Baxter. Energy-efficient neuronal computation via quantal synaptic failures. Journal of Neuroscience, 22 (11):4746–4755, 2002.

[55] B. Sengupta, M. B. Stemmler, and K. J. Friston. Information and efficiency in the nervous system—a synthesis. PLoS Comput Biol, 9(7):e1003157, 2013.

[56] M.-J. Cabirol-Pol, D. Combes, V. S. Fénelon, J. Simmers, and P. Meyrand. Rare and spatially segregated release sites mediate a synaptic interaction between two identified network neurons. Journal of neurobiology, 50(2):150–163, 2002.

[57] D. J. Schulz, J.-M. Goaillard, and E. Marder. Variable channel expression in identified single and electrically coupled neurons in different animals. Nature neuroscience, 9(3):356–362, 2006.

[58] A. Tejero-Cantero, J. Boelts, M. Deistler, J.-M. Lueckmann, C. Durkan, P. J. Gonçalves, D. S. Greenberg, and J. H. Macke. Sbi-a toolkit for simulation-based inference. arXiv preprint arXiv:2007.09114, 2020.

[59] L. Abbott and E. Marder. Modeling small networks, 1998.

[60] C. Durkan, A. Bekasov, I. Murray, and G. Papamakarios. Neural spline flows. In Advances in Neural Information Processing Systems, pages 7511–7522, 2019.

[61] F. Pedregosa, G. Varoquaux, A. Gramfort, V. Michel, B. Thirion, O. Grisel, M. Blondel, P. Prettenhofer, R. Weiss, V. Dubourg, J. Vanderplas, A. Passos, D. Cournapeau, M. Brucher, M. Perrot, and E. Duchesnay. Scikit-learn: Machine learning in Python. Journal of Machine Learning Research, 12:2825–2830, 2011.

[62] K. He, X. Zhang, S. Ren, and J. Sun. Deep residual learning for image recognition. In Proceedings of the IEEE conference on computer vision and pattern recognition, pages 770–778, 2016.

[63] N. Srivastava, G. Hinton, A. Krizhevsky, I. Sutskever, and R. Salakhutdinov. Dropout: a simple way to prevent neural networks from overfitting. The journal of machine learning research, 15(1):1929–1958, 2014.

[64] P. G. Constantine. Active subspaces: Emerging ideas for dimension reduction in parameter studies. SIAM, 2015.

[65] R. M. Neal. Slice sampling. The annals of statistics, 31(3):705–767, 2003.

